# A computationally efficient Bayesian Seemingly Unrelated Regressions model for high-dimensional Quantitative Trait Loci discovery

**DOI:** 10.1101/467019

**Authors:** L. Bottolo, M. Banterle, S. Richardson, M. Ala-Korpela, M-R. Järvelin, A. Lewin

**Affiliations:** Department of Medical Genetics, Cambridge, UK; The Alan Turing Institute, London, UK; MRC Biostatistics Unit, Cambridge, UK; Department of Medical Statistics, London School of Hygiene and Tropical Medicine, London, UK; Computational Medicine, University of Oulu and Biocenter Oulu, Oulu, Finland; NMR Metabolomics Laboratory, University of Eastern Finland, Kuopio, Finland; Center for Life Course Health Research, University of Oulu, Oulu, Finland; Biocenter Oulu, University of Oulu, Oulu, Finland; Department of Epidemiology and Biostatistics, Imperial College London, London, UK; MRC-PHE Centre for Environment and Health, Imperial College London, London, UK; Department of Life Sciences, Brunel University London, Uxbridge, UK

**Keywords:** Bayesian computation, Covariance reparametrisation, Graphical models, Markov Chain Monte Carlo, Metabolomics, Quantitative Trait Loci

## Abstract

**Motivation:** Our work is motivated by the search for metabolite Quantitative Trait Loci (QTL) in a cohort of more than 5,000 people. There are 158 metabolites measured by NMR spectroscopy measured in the 31-year follow-up of the Northern Finland Birth Cohort 1966 (NFBC66). These metabolites, as with many multivariate phenotypes produced by high-throughput biomarker technology, exhibit strong correlation structures. Existing approaches for combining such data with genetic variants for multivariate QTL analysis generally ignore phenotypic correlations or make restrictive assumptions about the associations between phenotypes and genetic loci.

**Results:** We present a computationally efficient Bayesian Seemingly Unrelated Regressions (SUR) model for high-dimensional data, with cell-sparse variable selection and sparse graphical structure for covariance selection. Cell-sparsity allows different phenotype responses to be associated with different genetic predictors and the graphical structure is used to represent the conditional dependencies between phenotype variables. To achieve feasible computation of the large model space, we exploit a factorisation of the covariance matrix. Applying the model to the NFBC66 data with 9,000 directly-genotyped Single Nucleotide Polymorphisms, we are able to simultaneously estimate genotype-phenotype associations and the residual dependence structure amongst the metabolites at the same time.

**Availability and implementation:** The R package `BayesSUR` with full documentation is available at https://cran.r-project.org/web/packages/BayesSUR/

## 1 Introduction

Integrating high-dimensional molecular biomarker data sets is a fundamental problem in genetic epidemiology and bioinformatics, in the search for molecular mediating mechanisms of the effects of genetic variants on clinical phenotypes. An important part of the analysis problem is to find associations between a set of genetic variants and downstream molecular phenotypes such as gene expression, proteomics, metabolomics or epigenetic data (Quantitative Trait Loci or QTL analysis). In the simplest analysis, univariate regressions are performed for each phenotype-genotype pair, needing post-hoc adjustment for multiple comparisons and ignoring any correlations between genotypes and between phenotypes. This is unlikely to be the best strategy for data where latent structures induce high levels of correlation between phenotypes, for example serum metabolomic profiles (Soininen et al., 2009; Kettunen et al., 2012), imaging and gene expressions. Comparison studies show that using multivariate quantitative phenotypes increases the statistical power in association tests compared to univariate analysis (Fusi et al., 2012; Inouye et al., 2012).

In this paper we develop a model designed for integrated multivariate QTL analysis, particular aimed at highly correlated molecular phenotypes. Our case study is in metabolomics Quantitative Trait Locus (mQTL), a powerful approach used to identify genes associated with metabolic markers of diseases, where the multivariate response is generally on the order of hundreds of metabolites. The model is also suitable for other multivariate molecular phenotypes. We have a dataset from the Northern Finland Birth Cohort 1966 (NFBC66) 31-year follow-up, consisting of 158 nuclear magnetic resonance (NMR) spectroscopy measured metabolites and over 9,000 directly genotyped Single Nucleotide Polymorphisms (SNPs) on chromosome 16, with a sample size of more than 5, 000 people. The metabolites set comprises lipoprotein particle concentrations, low molecular weight metabolites such as amino acids, 3-hydroxybutyrate and creatinine and different serum lipids, including free and esterified cholesterol, sphingomyelin and fatty acid saturation. These data exhibit strong residual correlation (Kettunen et al., 2012; Marttinen et al., 2014), even after accounting for the variance explained by all reported SNPs.

Brown et al. (1998, 2002) developed a general framework for Bayesian Variable Selection (BVS) models for multivariate outcomes (*s* responses) and multiple predictors (*p* covariates). In applications of this model, two alternative simplifying assumptions have been made in order to improve computational efficiency. The first simplifying assumption is to assume row-sparsity, in which a single set of predictors is selected across all response variables. Petretto et al. (2010) use this assumption for eQTL (expression Quantitative Trait Loci) analysis, with a dense residual covariance matrix across responses. Bhadra and Mallick (2013) also assume row-sparsity, but include sparse residual covariance selection between regressions, using graphical modelling based on decomposable graphs. In these models, row-sparsity induces conjugacy for regression coefficients and residual covariances, so these parameters are integrated out, resulting in a model search over the space of variable selection indicator variables. This approach has been used for small numbers of response variables (e.g., s around 10), with the number of predictors p in the thousands.

The second simplifying assumption is to assume conditional independence between regressions. Jia and Xu (2007); Bottolo et al. (2011); Scott-Boyer et al. (2012); Ruffieux et al. (2017, 2020a,b) take this approach for eQTL analysis, using hierarchical sparsity priors on the probabilities of associations, to share information across the gene expression outcomes. The assumption of residual independence between regressions ensures conjugacy of the regression coefficients and residual variances, enabling exploration of the full posterior space of selection models. In this scenario different predictors can be associated with different response variables without losing conjugacy. We refer to this as cell-sparsity. These hierarchical models can be used for larger numbers response variables, e.g., s in the hundreds or thousands.

Biologically complex phenotypes such as metabolomics and proteomics show strong correlation structures in comparison with transcriptomics data used in eQTL analyses, for which most BVS models have been developed. Therefore accounting for correlation to better reveal associations with genetic profiles, but also to infer their dependence structure, is becoming increasingly important to improve our understanding of integrated functioning of living organisms (Cichonska et al., 2016; Rodriguez-Martinez et al., 2017). Suitable statistical tools which can tackle these problems while retaining computational feasibility are thus necessary.

We present a Bayesian model for multivariate QTL analysis with correlated phenotypes, allowing for (i) cell-sparsity in the genotype-phenotype associations and (ii) residual dependence amongst phenotypes. We present two versions of the model, one with dense residual dependence structure and one with sparse covariance selection. Each instance of the multivariate regression model has the form of a Seemingly Unrelated Regressions (SUR) model. We exploit a factorisation of the covariance matrix parameter to enable faster computation using Markov Chain Monte Carlo (MCMC) methods. We are able to infer associations with thousands of candidate predictors (p over 9,000) multivariately on all responses. In the case of sparse covariance, thanks to the computationally efficient junction tree representation of a decomposable graph as its state variable (Green and Thomas, 2013), we also infer an underlying graph detailing the conditional independence relationships between hundreds of responses (*s* = 158).

The Bayesian framework has the advantage that it allows flexible sparsity priors on the regression and covariance coefficients, whilst being transparent about the overall level of sparsity. It also provides a rich output, full posterior distributions of coefficients and probabilities of variable inclusion.

The factorisation we use is related to the Cholesky decomposition of the precision matrix and was first proposed in the graphical modelling literature by Wermuth (1980) for multivariate Gaussian data with zero means and has been used in these so called “covariance selection” models for computational convenience and for interpretability of the now unconstrained transformed off-diagonal elements of the covariance matrix, see for example Pourahmadi (1999); Stingo and Marchetti (2014). Modelling Cholesky factors is also popular in econometrics in both conditional and simultaneous autoregressive models (Datta et al., 2019). Wang et al. (2012) use a similar idea in the context of sampling the parameters for the Bayesian lasso, and Zellner and Ando (2010) used this reparametrisation for a SUR model with fixed covariates, i.e., not resulting from variable selection.

To our knowledge this is the first model proposed for fully Bayesian analysis for multivariate QTLs allowing for both cell-sparsity and residual dependence. A similar model was proposed by Wang (2010) for autoregressive models in econometrics, with two computational algorithms, an MCMC algorithm using Gibbs updates (George and McCulloch, 1993) for variable selection called “indirect” and a “direct” algorithm involving numerical approximation of the marginal likelihood (Chib, 1995) combined with a Metropolis-Hastings algorithm for posterior models exploration. Both approaches work well for small data sets but are computationally prohibitive in high-dimensional space comprising hundreds of responses and predictors such as in molecular QTL work. Our novel contribution to the computational method is that we derive the priors (dense and sparse versions) in the space of the transformed covariance matrix and therefore run the whole computation in this space, enabling parallelisation over the response variables. We are thus able to estimate models for mQTLs with hundreds of response variables and thousands of predictors.

Section 2 introduces the model including the derived priors in the transformed space. Section 3 provides the posterior computations for the parameters and details the MCMC algorithm used to estimate them. In Section 4 the method is validated in an extensive simulation study, where we show that we can estimate larger models with more accuracy in considerably less computational time than the Wang (2010) software. We also obtain similar sensitivity but fewer false positives than the penalized likelihood method with simultaneous estimation of regression coefficients and covariance structure proposed by Rothman et al. (2010). Section 5 presents the results on the NFBC66 mQTL dataset, including visualisation both of the genotype-phenotype associations and of the residual dependence structure between metabolites.

Further details of model derivations and posterior updates are available in the Supplementary Material. An R package BayesSUR with full documentation is available at https://cran.r-project.org/web/packages/BayesSUR/.

## 2 Model

The model can be seen as a set of regressions for multivariate phenotype responses *Y* = (***y***_1_,⋯, ***y_s_***), ***y***_*k*_ = (*y*_1*k*_,⋯, *y_nk_*)^*T*^, for *k* = 1,…, *s* and corresponding covariate genotype matrices *X_k_* with dimensions *n* × *p*. We assume independence between samples, but allow for dependence across responses. Moreover, we assume that the same set of predictors are available for all responses. The same set of genotypes may be used for all regressions, but this is not necessary. The predictors may be continuous or categorical, hence the model accommodates the usual additive genetic association models using observed and imputed allele counts or can be extended to more complex genetic models including interactions.

Variable selection is performed on the predictors using binary indicators vector *γ_k_* = (*γ*_*k*1_,⋯, *γ_kp_*)^*T*^, where *γ_kj_* is 1 if covariate *j* is included in the regression for response *k* and 0 if not. We use the shorthand notation *X_γk_* for the columns of *X_k_* selected by the vector *γ_k_* and similar for *β_γk_*. Thus, we can write the set of linked regressions as

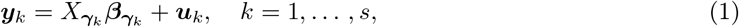

but most importantly the residuals will be correlated, i.e., *u_i_* = (*u*_*i*1_,⋯,*u_is_*) ~ N(**0**, *C*).

We can also write the likelihood for this model as

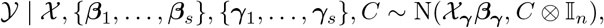

where 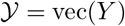, vec(·) being the vectorisation operator, *γ* = vec(*γ*_1_,⋯, *γ_s_*), *β_γ_* = vec(*β*_*γ*1_,⋯, *β*_*γs*_) and *X_γ_* is a block-diagonal matrix with *X_γk_* as the *k*-th diagonal element.

Following George and McCulloch (1993), we set *β_kj_* = 0 conditional on *γ_kj_* = 0, while non-zero coefficients follow a diffuse Normal distribution, i.e., 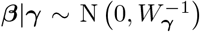. The precision matrix *W* is generally decomposed into a shrinking coefficient, say *w*, and a matrix that governs the covariance structure. Here, we use 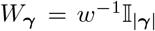, with an Inverse-Gamma prior on *w* and |*γ*| the number of selected predictors across all outcomes. A number of sparsity inducing priors for *γ* have been used in the literature, the most common being *γ_kj_* ~ Ber(*π*) with a fixed or random *π*. Another choice in QTL analysis is the “hotspot detection” prior used in Bottolo et al. (2011), which decomposes the inclusion probability into an overall sparsity level for each outcome and a propensity parameter for each predictor, i.e., *γ_kj_* ∼ Ber(*o_k_* × *π_j_*). In this work, we use the hotspot detection model with a Beta prior on *o_k_* and a Gamma prior on the propensity *π_j_* and its simplified version with *π_j_* = 1, *j* = 1,⋯,*p*, which corresponds to a Beta-Binomial sparsity prior for each response.

### 2.1 Factorisation of the likelihood

If one assumes either a diagonal *C* or row-sparsity for *γ_k_*, with conjugate priors on *C* and *β_γ_*, both *β_γ_* and *C* can be integrated out analytically (Bhadra and Mallick, 2013). In our model the usual priors on these parameters lose conjugacy and can’t be integrated out. Nonetheless, the full conditionals retain their simple forms, so it is straightforward to write a Gibbs sampler for the posterior distributions (Holmes et al., 2002). The computational time needed is however prohibitive for most high-dimensional settings.

To overcome this issue, we decompose the covariance matrix *C* iteratively as

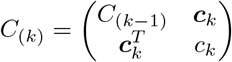

for all *k* = 2,…, *s*, with *C*_(*s*)_ = *C* and *C*_(1)_ = *c*_1_ = *C*_11_ (the scalar variance of response 1) and *c*_1_ is null. Thus, each *C*_(*k*)_ is the marginal covariance matrix for responses 1,…,*k*, *c_k_* is the variance of response *k* and *c_k_* is the vector of covariances between response *k* and responses {1,…, *k* – 1}.

With this decomposition, the likelihood can be factorised (e.g., Giri (2014)) as

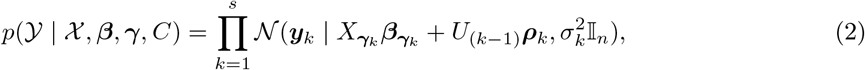

where *U*_(*k*−1_) = *Y*_(*k*–1)_ – (*X*_*γ*1_ *β*_*γ*1_ *X*_*γ*2_ *β*_*γ*2_ ⋯ *X*_*γk*−1_ *β*_*γk*−1_) is a matrix consisting of the first *k* – 1 residuals from the original linked regressions where *U*_(0)_ is null. For *k* = 1 the likelihood simplifies to 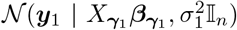. The parameters 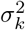 and *ρ_k_* are also defined through the reparametrisation of the residual covariance matrix, i.e., 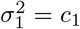 and

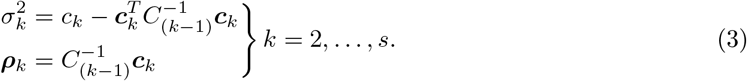

Note that the joint distribution 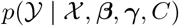 is the same regardless of the order used for the decomposition since we are simply factorising it by chain-conditioning. From Equation (2) is straightforward to see that 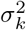 is the residual variance of response *k* conditioned on *U*_(*k*-1)_ and *ρ_k_* is the (*k* – 1)-vector of regression coefficients on the same *U*_(*k*-1)_ residuals. The likelihood is thus decomposed into a product of independent (conditionally on the new parameters) factors over the outcomes. The novelty of our approach is two-fold. First, we estimate our model completely in the reparametrised space, deriving priors and posterior full conditionals for the parameters 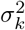 and *ρ_k_*. Second, this allows us to update these parameters in parallel, which greatly increases the computational efficiency of this model.

### 2.2 Prior for dense residual dependence

n order to take advantage of the factorisation of the model across responses, we must also transform the model priors. For modelling dense dependence structure between responses, we use an Inverse-Wishart prior on the original covariance matrix *C* ~ IW(*ν, M*). We use standard matrix properties of the Inverse-Wishart distribution (Dempster, 1969; Dawid, 1981; Roverato, 2000) to calculate the transformed prior. The *C*_(*k*)_ is a submatrix of *C*, thus it also has an Inverse-Wishart distribution. The new parameters are related to the block structure of the inverse of this matrix, 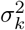 being the Schur complement of *C*_(*k*-1)_ in *C*_(*k*)_ and 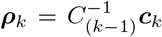. Decomposing *M* conformally with *C* into 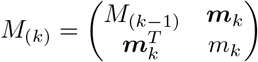 for *k* = 2, …, *s*, the priors on the changed variables become

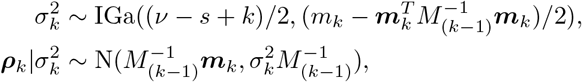

for *k* = 2,…, *s* and 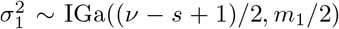. From the prior on 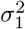, we can see that we the degrees of freedom in the Inverse-Wishart distribution must be chosen to be *ν* > *s* – 1. Moreover, from these equations, we can see that we obtain independent priors for 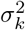 and *ρ_k_* for different *k*. Thus, since both likelihood and prior factorise across responses, the posterior full conditionals also factorise and hence the MCMC update of the residual covariance parameters can be parallelised.

### 2.3 Prior for sparse residual dependence

To model sparsity in the residual dependency structure, we introduce a decomposable graph *G* such that variables are conditionally independent if there is no direct edge between them in the graph (Lauritzen, 1996). Conditional on the graph, we use the Hyper Inverse-Wishart prior on the original covariance matrix *C* ~ HIW_*G*_(*v, M*) (Carvalho et al., 2007). This is defined as the distribution such that the covariance matrix for each prime component in the decomposable graph is marginally Inverse Wishart, i.e., *C*_P_q__ ~ IW(*v* – (*s* – |*P_q_*|),*M_P_q__*), where *P_q_* is the *q*-th prime component, *M_P_q__* is the sub-matrix of *M* corresponding to *P_q_* and |*P_q_*| is its cardinality.

In the following, we derive the corresponding prior for the transformed variables 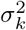 and *ρ_k_* under the assumption of a sparse covariance matrix. A more in depth derivation of the following results can be found in the Supplementary Material Section S.1.1. Briefly, the decomposability of the graph G allows us to define a sequence of complete, overlapping, subgraphs (i.e., cliques) called “prime components” *P_q_, q* ∈ (1,…, *Q*) that can be ordered in such a way that for every *q* > 1 there exists *m* < *q* such that *P_q_* ⋂ *H_q_* ⊂ *P_m_*, where 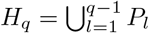 for *q* = 2,…, *Q*. We also define the separators *S_q_* = *P_q_* ⋂ *H_q_* and residuals *R_q_* = *P_q_* \ *S_q_* for *q* = 2,…, *Q*. The nodes of G can also be arranged according to a so-called “perfect elimination ordering”, denoted by *ξ*, which implies that if Λ_*ξ*_*k*_*ξ*_*l*__ = 0 then *ρ*_*ξ*_*k*_*ξ*_*l*__ = 0, where Λ is the precision matrix *C*^-1^ (Paulsen et al., 1989).

Due to this correspondence between *Λ*_*ξ*_*k*_*ξ*_*l*__ = 0 and *ρ*_*ξ*_*k*_*ξ*_*l*__ = 0, the transformation in Equation (3) decomposes across the prime components, so that, for nodes ordered according to a perfect elimination ordering, the new variables 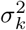 and *ρ_k_* are defined within the prime components, i.e.,

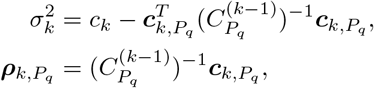

where 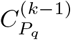 is the sub-matrix of *C_P_q__* with variable *k* removed and *c_k,P_q__* is the final column of C_P_q__ without the last element. All other elements of *ρ*_*k*_ are zero.

To write the densities explicitly, we need to order the nodes using the perfect elimination order *ξ*, which respects the perfect ordering of the prime components. With this ordering, we find that the Hyper Inverse-Wishart prior on *C* is transformed to

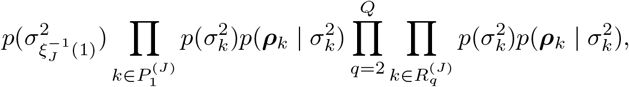

where the ordering of nodes within each residual does not matter. The corresponding prior densities are

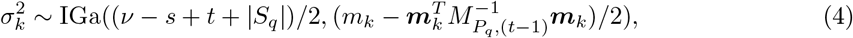

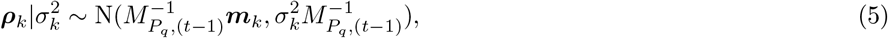

where *q* is the index of the prime residual that node *k* belongs to in the particular node ordering of the graph, for all *k*, except for the first node, i.e., 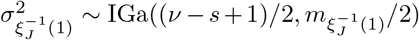. The sets *P_q_* and *S_q_* are defined above. The index *t* in Equations (4) and (5) is the index of the node within the graph residual component and is given by *ξ_k_* – |*H_q_*|. Here, we have applied similar arguments as for the dense covariance case presented in Section 2.2, using the properties of block matrices and the Inverse-Wishart distribution. Again, we see that the priors factorise over responses, so the posterior full conditionals in Equations (4) and (5) also factorise enabling faster computation of the model through parallelization of the MCMC updates. An important feature of working with this transformation is that we do not need to do any completion operation to fill in the covariances between the separated parts of prime components (Carvalho et al., 2007) since these correspond to the zeros in the *ρ_k_, k* = 1,…, *s*, parameters.

We parametrise *G* using the junction tree representation of a decomposable graph as its state variable proposed by Green and Thomas (2013). A decomposable graph may have many junction tree representations. However, for a given junction tree, the implied graph is uniquely determined. We use a prior on the junction tree *J* which is proportional to a Binomial distribution on the number of edges |*J*| in the graph, i.e., 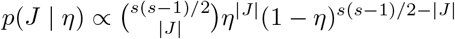. Since sparser graphs in general have more junction tree representations (Thomas and Green, 2009), this prior favours sparse structures. Finally, we use a conjugate Beta prior on the hyper-parameter *η*.

### 2.4 Summary of full model

In this section, we summarise the full model with all its conditional dependencies. We provide the version using the sparse covariance structure. The dense covariance case is as below, except that there are no *J* or *η* parameters and the distributions for 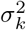 and *ρ_k_* are the simpler expressions presented in Section 2.2. In Sections 2.2 and 2.3 we have illustrated the results in terms of a general matrix *M* in the (Hyper) Inverse-Wishart distribution. In our implementation we use 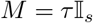.

The joint distribution is

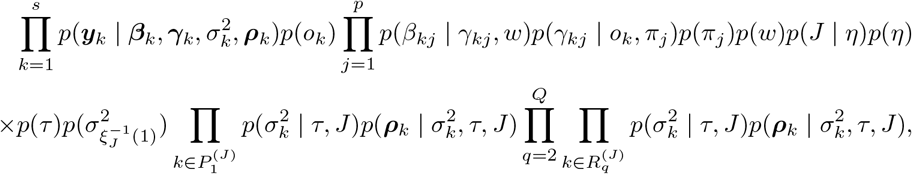

where

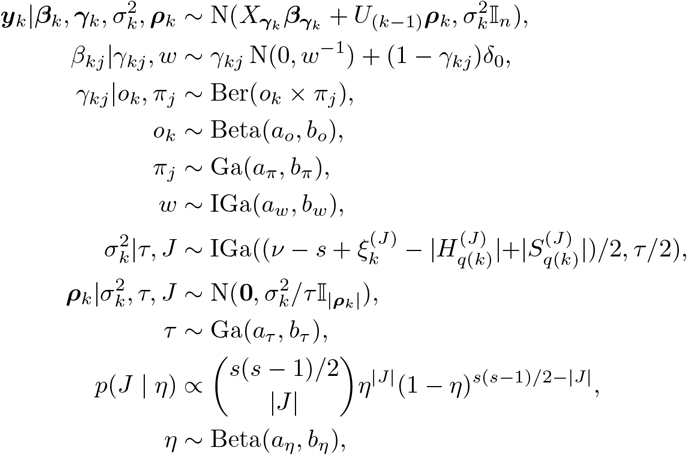

with *U*_(*k*−1)_ is defined as in Equation (2) and *δ*_0_ is the Dirach delta function in 0. The parameter *J* stands for the junction tree representing the graph and |*J*| is the number of edges in the graph represented by the junction tree. *ξ*^(*J*)^ is a perfect elimination ordering of the nodes in the graph. Both 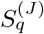 and 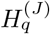 depend on the graph and are defined in Section 2.3. The index *q*(*k*) is the index of the prime residual 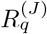 that node *k* belongs to in the current node ordering for graph *J*.

Finally, the parameters *a_o_, b_o_, a_π_, b_π_, a_w_, b_w_, a_τ_, b_τ_* and *a_η_, b_η_* are fixed. The degrees of freedom ν > s — 1 in the Inverse-Wishart distribution is also fixed.

## 3 Posterior computations

In the original model space, posterior full conditionals for *β_γ_* and *C* are available analytically (Holmes et al., 2002). However, these updates require inverting at every MCMC update both the |*β_γ_*|×|*γ*| quadratic form in the selected columns of the design matrix *X_γ_* and the *s* × *s* matrix for the covariance matrix *C*. Additionally, the update of *γ* and all other unknowns where the likelihood is involved, require the heavy computation of the non-factorised likelihood. Our approach, based on the reparametrisation of *C* which leads to the factorisation of the model and the introduction of a sparse precision matrix *via* the junction tree representation of the decomposable graph *G* as its state variable, allows us to introduce a much more computational efficient MCMC scheme that scales well in high-dimensional settings.

Zellner and Ando (2010) used the same reparametrisation in a simpler SUR model without variable selection, using Jeffrey’s priors. They devised a Direct Monte Carlo procedure for *β*, *σ*^2^ and *ρ*. Their method uses an approximation to the full conditionals, with an additional resampling step for the *β*. However, it is possible to recover the correct posterior full conditional for *β*, avoiding unnecessary and computationally prohibitive resampling steps as we show below.

To sample from the posterior distribution of the binary indicators vector *γ*, we use the Evolutionary Stochastic Search (ESS) algorithm (Bottolo and Richardson, 2010; Bottolo et al., 2011; Lewin et al., 2016), which uses a particular form of Evolutionary Monte Carlo as defined in Liang and Wong (2000). Within this framework, posterior samples of *β_γ_*, *σ*^2^ and *ρ* are obtained by employing a Gibbs sampler, but used instead in the joint updates of {*γ*, *β_γ_*} and {*J*, *σ*^2^, *ρ*}. Specifically, the posterior full conditionals for *β_γ_* and *σ*^2^, *ρ* are used as proposal distributions in the joint updates with *γ* and *J*, respectively, since it reduces the posterior correlation between *γ* - *β_γ_* and *J* – {*σ*^2^, *ρ*}. In this set-up, the proposal and target densities cancel out in the Metropolis-Hastings acceptance ratios. This is known as “implicit marginalisation” (Holmes and Held, 2006; Alexopoulos and Bottolo, 2020) since the resulting acceptance ratio does not contain the current and proposed values of *β_γ_* or {*σ*^2^, *ρ*} and it has been shown to greatly improve mixing of the structural parameters *γ* and J which are in our case the main focus of inference.

We have derived the full conditionals for the regression parameters and state the results here. Further details regarding the derivations can be found in the Supplementary Material Section S.1.5. The posterior conditional for the non-zero regression coefficients is

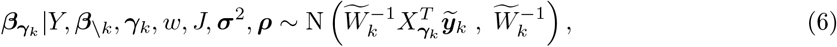

where the subscript “\*k*” implies that the vector of the regression coefficients consists of all the elements except those that are related to the *k*-th response,

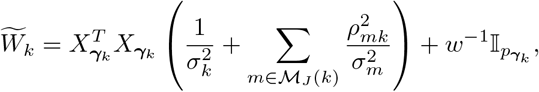

and

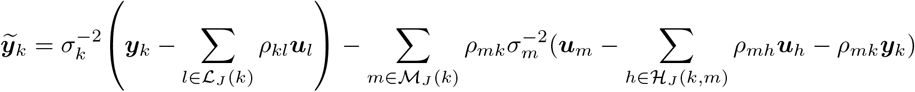

with *u_k_* defined as the residuals given in Equation (1).

In the sparse covariance case, the index sets are defined with respect to the perfect elimination order *ξ*, i.e.,

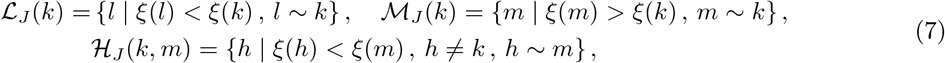

where *l* ~ *k* means that nodes *l* and *k* are in the same prime component. In the dense case, these reduce to

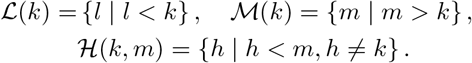

The posterior updates of the reparametrised covariance parameters depend on the ordering of the nodes and prime residuals of the graph

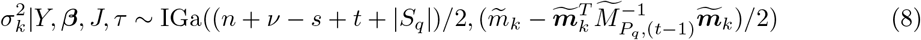

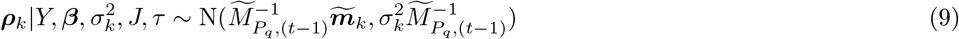

with 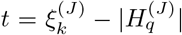 as before, 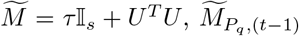 and 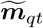 are sub-matrices of 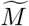 defined conformally with previous transformations. In the dense covariance case, the posterior updates reduce to the following equations (with any ordering on the outcomes)

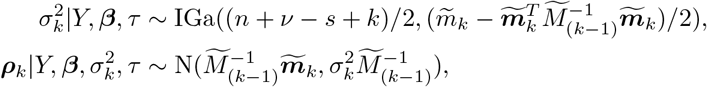

To efficiently explore the graphical structure *G*, we use the sampler introduced by Green and Thomas (2013), making use of the junction tree representation of a decomposable graph as its state variable to allow for bolder, multi-edge proposals in the graph space. The edge probability *η* is updated via a Gibbs perturbation. All other unknowns are updated via Metropolis-within-Gibbs updates with adaptive proposal distributions (Roberts and Rosenthal, 2009). Algorithm 1 provides an overview of the designed MCMC algorithm to sample from the joint posterior distribution *p*(*β*, *γ*, *σ*^2^, *ρ*, *J*, *π*, *o*, *w*, *τ*, *η*|*Y*).

Note that, even though each sample is from a decomposable model, the sampler allows us to discover non-decomposable graph structures via Bayesian Model Averaging of the marginal edge inclusion probabilities. As the graph is updated, the perfect elimination ordering *ξ* changes, hence we do not retain the sampled values of *σ*^2^ and *ρ*. Moreover, as we are interested mainly in structure learning, both for variable and covariance selection, we consider the reparametrised covariance as nuisance parameters.

## 4 Simulation study

We evaluate the performance of the reparametrised multivariate sparse SUR model and our efficient sampler in simulated mQTL data. We first investigate the effect of allowing for residual dependence in the phenotypes, by comparing dependent and independent covariances within our own model. We then compare our model with sparse dependence structure against other methods that also allow for dependence. For all the work in the simulation study, we employ the same priors as we use for the mQTL analysis of the NFBC cohort data, see Section 5 for details, except in the comparison with other models that allow covariance selection where we use the simplified version of the hotspot detection prior with *π_j_* = 1, *j* = 1,…,*p*.

**Algorithm 1.**
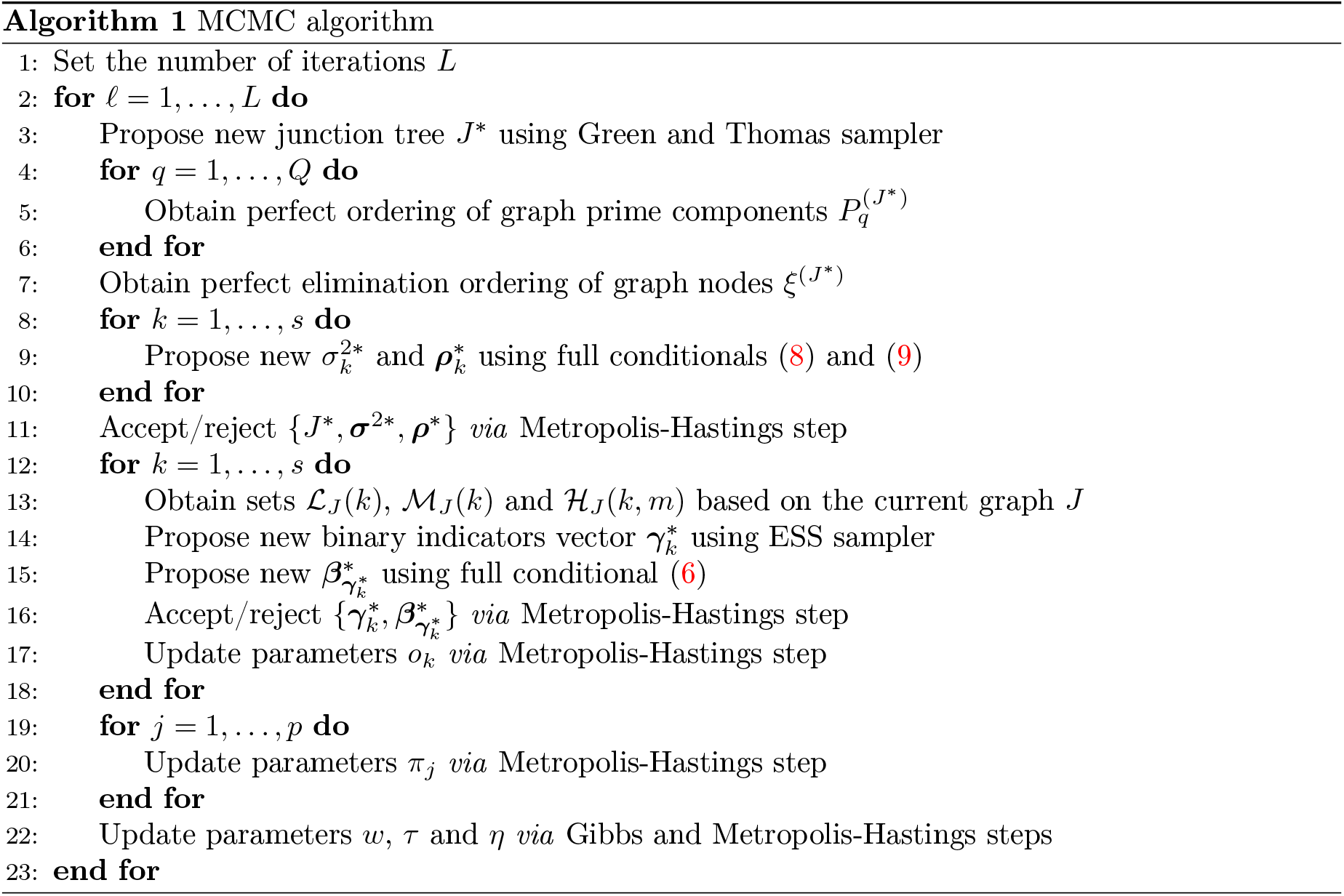
MCMC algorithm

### 4.1 Comparison with models without covariance selection

We validate our method against the Hierarchical Evolutionary Stochastic Search (HESS) algorithm of Bottolo et al. (2011) in a synthetic setting. Following Richardson et al. (2010) and Bhadra and Mallick (2013), we set up our simulation study by randomly subsampling *p* = 300 SNPs from our real *-omics* dataset (see Section 5). This forms our covariate set *X* and allows us to mimic real correlation effects and linkage disequilibrium between genetic markers that would be difficult to simulate artificially. The observed correlations between predictors ranges from small to over 0.8 in absolute value. We set *n* = 200 and s = 30 and proceed by selecting the correlation structure for the outcomes in form of a graph. We explore three graphical structures, i.e., a block diagonal, a decomposable and a non-decomposable model.

To present a range of possible association patterns between outcomes and predictors, we fix (conditionally on the selected graph structure) the binary indicators vector *γ* so that different sets of predictors display associations with, i.e., *all* outcomes (representing true hotspots), all outcomes within each prime component, all outcomes within each residual component (so predictors are linked only with correlated outcomes and not to conditionally independent ones) and finally with a set of selected outcomes that spans multiple components, so that selected predictors are linked to both correlated and (conditionally) independent outcomes. See Supplementary Figure S.2 for an example of the generated structures.

With the structure fixed, we sample the non-zero regression coefficients from a N(5,1), so that most of them are distinct from zero, and the residuals from a matrix variate Normal distribution, i.e., MN (0,∥_*n*_,*C*). *C*^-1^ is sampled from a G-Wishart distribution, *W_G_* (*s* + 2, *M*), using the R package BDgraph (Mohammadi and Wit, 2019) with *M* = *αR, R* a correlation matrix with *r* on the off-diagonal elements and *α* ∈ ℝ^+^ to obtain data sets with the desired noise.

We consider two summaries of signal to noise, designed to be sensitive to the information contained in the predictors and in the covariance matrix respectively:

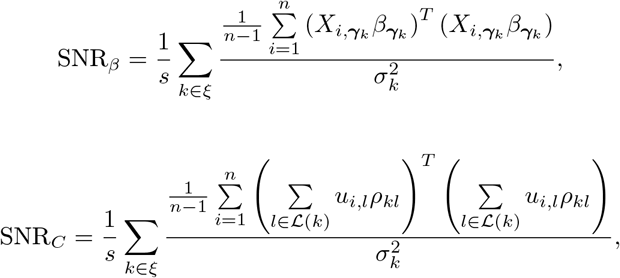

where *σ*^2^ and *ρ* are the reparametrised values of the covariance matrix *C* and 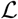 is as defined in Equation (7). We observe that SNR_*C*_ is highly correlated with the off-diagonal (residual) correlation *r*. Thus, we parametrise the simulation study in terms of *G* (block-diagonal, decomposable and non-decomposable), *r* ∈ {0.3,0.6,0.9} and SNR_*β*_. For each value of *G* and *r*, we generate multiple data sets with different *α* values and use the data with resulting SNR_*β*_ within 10% of each of the desired values of 5, 15 and 25. Based on this criteria, we simulate 20 replicates for each combination of the parameters and run both sparse BayesSUR model and HESS for 250,000 iterations of which 50,000 as burn-in.

To report on performance, we focus on posterior marginal inclusion probabilities, i.e., the average over the MCMC iterations of *γ_kj_*, *k* = 1,…,*s*, *j* = 1,…,*p*. Figure 1 shows the average ROC curves over 20 replicates for each simulation set-up corresponding to SNR_*β*_ = 5, the lowest signal-to-noise ratio, for both BayesSUR with covariance selection and HESS. The results correspond to our expectations, i.e., HESS is known to perform relatively well even in cases where residuals are correlated (Bottolo et al., 2011; Lewin et al., 2016) as long as *r* is not too high. In most cases though, especially at higher r, HESS estimates are more noisy and more false positives are picked up due to the confounding effect of the correlations. BayesSUR has a more marked separations between true and false positive signals and overall returns less noisy estimates (see Supplementary Figure S.5, top panels). The ROC curves relative to the other SNR_*β*_ levels are reported in Supplementary Figures S.3 and S.4.

**Figure 1:**
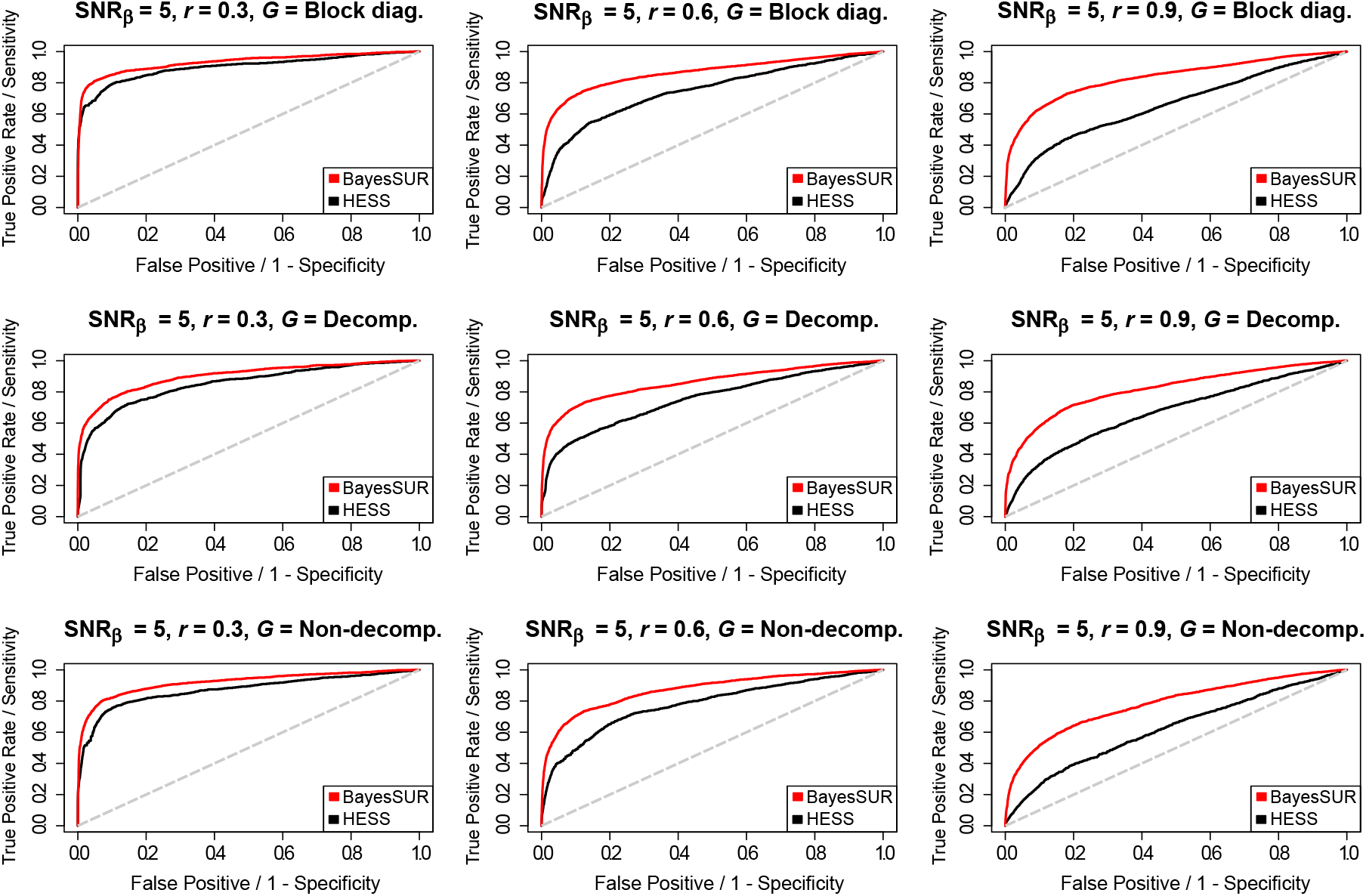
Averaged (over 20 simulated replicates) ROC curves for BayesSUR with covariance selection (red line) and HESS (black line) to compare the variable selection performance of the two methods for different combinations of the simulated graphical model G and the (residual) correlation between responses *r* and with signal-to-noise ratio for the predictors SNR_*β*_ = 5.

BayesSUR is also able to recover simultaneously the conditional (in)dependence structure of the residuals. Table 1 shows the average over 20 simulated replicates of true positive rates (TPR) and false positive rates (FPR) for graph edges found by thresholding Pr(*G_kk′_* = 1 | data) at 0.5 probability level. The graphs are in general well estimated. For non-decomposable graphs, there is a tendency toward over-inclusion, as we would expect based on Fitch et al. (2014), who find that graphs constrained to be decomposable converge to a close (in the graph space), more dense, chordal graph alternative (see for example Figure S.5 in the Supplementary Material). The estimated TPR and FPR for different values of SNR_*β*_ do not differ significantly from the one presented here. See, for details, Supplementary Table S.1.

**Table 1:** Averaged (over 20 simulated replicates) true positive and false positive rates for the graph estimation after thresholding at 0.5 the posterior marginal edge inclusion probabilities. Results are reported for SNR_*β*_ = 5 and for different combinations of the graphical model *G* and the (residual) correlation between responses *r*.

| $G$ | $r$ | True positive rate | False positive rate |
| --- | --- | --- | --- |
| Block diagonal | 0.3 | 0.937 | 0.008 |
| Block diagonal | 0.6 | 0.960 | 0.036 |
| Block diagonal | 0.9 | 0.953 | 0.016 |
| Decomposable | 0.3 | 0.935 | 0.085 |
| Decomposable | 0.6 | 0.958 | 0.108 |
| Decomposable | 0.9 | 0.922 | 0.127 |
| Non-decomposable | 0.3 | 0.962 | 0.027 |
| Non-decomposable | 0.6 | 0.986 | 0.117 |
| Non-decomposable | 0.9 | 0.979 | 0.146 |

### 4.2 Comparison with alternative covariance models

We compare the performance of BayesSUR to two different software implementations of a sparse seemingly unrelated regressions (SSUR) model by Wang (2010). SSUR Indirect performs posterior computation of the SSUR model using MCMC, where the regression coefficients are sampled using the Gibbs sampler described in George and McCulloch (1993). SSUR Direct uses the marginal likelihood approach of Chib (1995) for “direct” variable selection of important predictors and non-zero entries of the sparse inverse covariance matrix *via* Metropolis-Hastings steps. The Matlab version of the SSUR code is available from the author web site. For the “indirect” version, we run the algorithm for 5×10^5^ iterations with 10^5^ as burn-in, storing the MCMC output every 500 iterations. For the “direct” version, we run the algorithm for 2×10^3^ iterations with 10^3^ as burn-in. In each iteration, the calculation of the marginal likelihood requires 500 extra samples from the Gibbs sampler, including 100 as burn-in. Overall, the algorithm is run for 10^6^ iterations. All hyper-parameters and proposal densities are left unchanged as originally set-up in the Matlab code. The prior probability of inclusion is set equal to 0.1 in both versions of the Matlab code. We run BayesSUR with covariance selection for 5×10^5^ iterations with 10^5^ as burn-in, two parallel chains in the EES sampler and matching the hyper-parameters of the Beta-Binomial prior on the inclusion probability with the prior used in the SSUR algorithms.

For each scenario, we simulate 20 replicates with n = 150, p = 30 and s = 20. Out of 30 × 20 regression coefficients, 120 (20%) are simulated from an uniform distribution in (–2,2). We selected at random the same proportion of cells in the 30 × 20 matrix of regression coefficients and assigned to them the simulated values, while the other cells are set to zero. In our experiment, for each response, on average between 2 and 10 non-zero regression coefficients are assigned. To generate the correlated predictors, we follow Rothman et al. (2010) and simulate, for each *i* = 1, …,*n* and *k* = 1,…,*s*, *x_ik_* ~ N(0, *V*), where 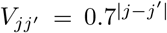 is the (*j, j’*)-th element of *V*, *j,j*’ = 1, …,*p*, implying the same unit marginal variance. The inverse error covariance *T*^-1^ is a Toeplitz matrix with value 0.5 in the first principal diagonal, 0.5 and 0.4 in the first two principal diagonals and 0.5, 0.4 and 0.3 in the first three principal diagonals in Scenario 1, 2 and 3, respectively. In all scenario considered, the sparse diagonal inverse error covariance is positive definite with 19 (10%), 37 (19%) and 54 (28%) non-zeros entries in Scenario 1, 2 and 3, respectively, while the corresponding covariance matrices are dense. Finally, the responses are generated from a Normal matrix variate distribution using the simulated matrix of regression coefficients, the predictors matrix and the dense covariance matrices, i.e., 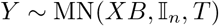, where *B* is the *p* × *q* matrix of the simulated regression coefficients and *T* is the inverse of the *q* × *q* Toeplitz matrix.

Figure 2 shows the ROC curves obtained from the simulation study distinguishing between the estimation of the non-zeros regression coefficients (top panels) and the estimation of non-zero entries of the inverse error covariance (bottom panels). From the plots, it is apparent that BayesSUR (with or without covariance selection) has better or similar performance to SSUR. It is more efficient than SSUR Direct in all scenarios considered due to the expensive computation of the approximate marginal likelihood that prevents running the algorithm for many iterations. BayesSUR performs better than SSUR Indirect whose performance deteriorates as the estimation of the sparse inverse error covariance becomes less sparse (Scenarios 2 and 3). A closer inspection of the MCMC output shows that in Scenarios 2 and 2 both versions of the SSUR algorithm incorrectly estimate that the responses are almost independent conditionally on the estimated regression coefficients (results not shown).

**Figure 2:**
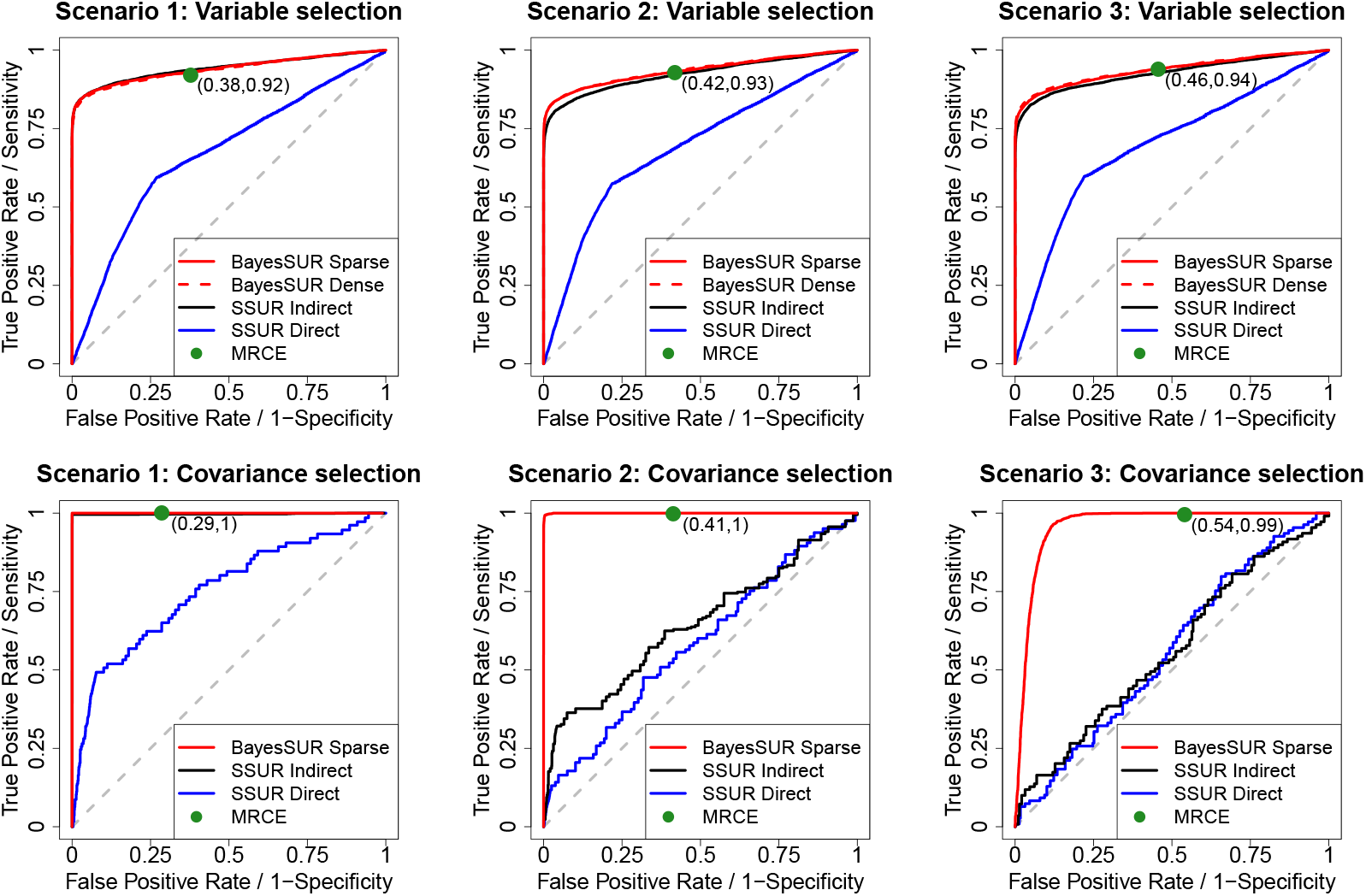
Averaged (over 20 simulated replicates) ROC curves to compare the selection performance of the non-zero regression coefficients (top panels) and non-zero elements of the precision matrix (bottom panels) for the methods considered: BayesSUR with covariance selection (solid red line), BayesSUR with dense covariance estimation (dashed red line), SSUR Indirect (black line), SSUR Direct (blue line) and MRCE (green dot). For MRCE, each dot represents the averaged specificity and sensitivity of the corresponding penalised likelihood solution. Different scenarios are obtained by specifying distinct Toeplitz matrices for the inverse error covariance.

We also compare the performance of BayesSUR with MRCE, the penalized likelihood method with simultaneous estimation of the regression coefficients and the covariance structure proposed by Rothman et al. (2010). While the power to detect non-zero regression coefficients and non-zero elements of the precision matrix is similar to BayesSUR, in all simulated scenarios, MRCE seems to include a larger number of false positives, in particular in the covariance selection.

In addition, we examine the computational time of the algorithms presented in this section. For the Bayesian algorithms, we match the values of the sparse priors hyper-parameters and, as far as possible, the total number of iterations. For MRCE, we select the option *cv*, i.e., the penalty parameters λ_1_ ∈ λ_1_ for the regression coefficients and λ_2_ ∈ λ_2_ for the covariance structure are chosen by using a 5-fold cross-validation procedure. We also specify different dimensions of the candidate penalty vectors λ_1_ and λ_2_ to check the impact of this choice on the computation time. All algorithms are run on an Intel(R) CPU 2.60GHz with 64 Gb memory.

Table 2 shows that BayesSUR is 20 times faster than SSUR Direct in all scenarios considered. Interestingly, it is also almost 10 times faster than SSUR Indirect with the SSVS Gibbs sampler. This is due to the effect of the direct manipulation of the junction tree representation of a decomposable graph (Green and Thomas, 2013) used in this work, in contrast to the computational expensive evaluation of the decomposability after edge perturbation employed in Wang (2010) and originally proposed by Giudici and Green (1999). The different computational efficiency depends also on the programming language used by the two algorithms, C++ and Matlab, respectively. Note that we employ a single core to run BayesSUR in order to make the comparison with other methods fair. However, large computational gains can be achieved by using a multi-core parallel computing architecture such as Message Passing Interface to exploit the parallelization of step 9 in Algorithm 1. The computational time of MRCE greatly depends on the number of candidate values where the 5-fold cross-validation procedure is performed. At the default value, 4 equally spaced grid points, the algorithm is very fast, but it becomes slower than BayesSUR when 200 candidate values are specified. Moreover, in contrast to BayesSUR, the sparser the graph, the slower MRCE becomes. Similarly to the number of iterations in MCMC algorithms, for penalised likelihood methods the choice of the number of candidate penalty values depends on the trade-off between accuracy and computational time.

**Table 2:**
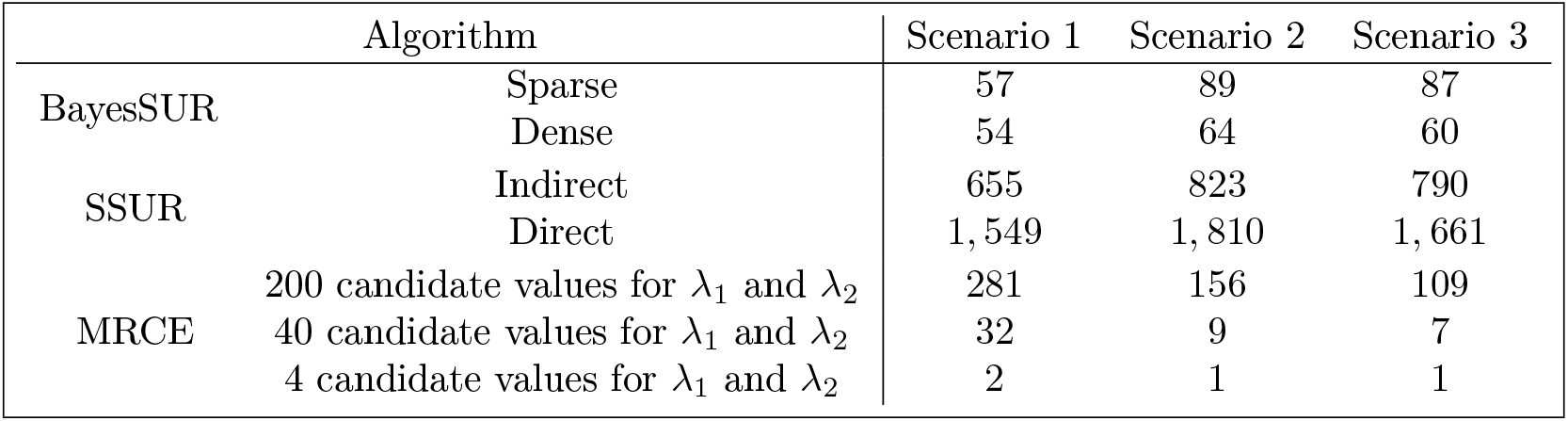
Averaged (over 20 simulated replicates) computational time in minutes for the algorithms considered: BayesSUR with covariance selection (Sparse), BayesSUR with dense covariance estimation (Dense), SSUR Indirect, SSUR Direct and MRCE with different numbers of candidate values for the penalty parameters where the cross-validation procedure is performed.

| Algorithm |  | Scenario 1 | Scenario 2 | Scenario 3 |
| --- | --- | --- | --- | --- |
| BayesSUR | Sparse | 57 | 89 | 87 |
|  | Dense | 54 | 64 | 60 |
| SSUR | Indirect | 655 | 823 | 790 |
|  | Direct | 1,549 | 1,810 | 1,661 |
| MRCE | 200 candidate values for $\lambda_1$ and $\lambda_2$ | 281 | 156 | 109 |
| | 40 candidate values for $\lambda_1$ and $\lambda_2$ | 32 | 9 | 7 |
| | 4 candidate values for $\lambda_1$ and $\lambda_2$ | 2 | 1 | 1 |

Finally, we repeat the same analysis presented in this section with s = 150 responses to mimic the number of responses of the real data example presented in the next section. Results are similar to those presented here, although the analysis becomes computationally prohibitive for both SSUR Direct and SUUR Indirect. Details of the selection performance of the different methods as well as their computational time are shown in Supplementary Figure S.6 and Table S.2, respectively.

## 5 Metabolite Quantitative Trait Loci (mQTL) analysis in the Northern Finnish Birth Cohort

In this section we present our results of the mQTL analysis of the NFBC66 data. The serum metabolic data are from the 31-year follow-up study of the NFBC66 and based on a widespread metabolomics platform in epidemiology and genetics (Würtz et al., 2017). After quality control and data cleaning, the data consist of *p* = 9, 310 directly genotyped SNPs on chromosome 16 and *s* = 158 metabolite concentrations, measured on *n* = 5, 154 individuals. The metabolites are normalised and standardised *via* the inverse rank-Normal transformation, following Kettunen et al. (2016).

Thanks to growing evidence in favour of pleiotropy (the association of multiple phenotypes with the same locus) in mQTL analysis (Sabatti et al., 2009), we expect these associations to be driven by a handful of SNPs that associate with numerous metabolites. To drive the variable selection procedure we will therefore use the *hotspot* prior introduced by Bottolo et al. (2011) which expresses the prior probabilities for variable inclusion into overall sparsity level *o_k_* for outcome k and a propensity parameter *π_j_* for each predictor *j* with *γ_kj_* ~ Ber (*o_k_* × *π_j_*), *π_j_* ~ Ga(1/2,1/2) (E(*π_j_*) = 1, Var(*π_j_*) = 4) and *o_k_* ~ Beta(*a_o_, b_o_*) under the constraint *o_k_* × *π_j_* ≤ 1 ∀_*j*_, *k*. The hyper-parameters *a_o_, b_o_* are chosen so that the average model size and its variance for each outcome is small, as we want to enforce a strong sparsity in the model, i.e., *E*(*o_k_*) = 2 and Var(*o_k_*) = 2. These values imply *a priori* a range of associations for each response between 0 and 6. We use independent N(0, w) priors on non-zero regression coefficients which correspond to 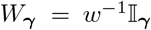 and let the prior matrix in the InverseWishart for the covariance be diagonal, i.e., 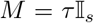. Since we standardise and centre all responses and predictors, the hyper-parameters for *τ* and *w* are set such that these variances are centred on small values but, at the same time, allowing the respective prior to be diffuse (*a_w_* = *b_w_* =0.1 and *a_τ_* = *b_τ_* = 0.1). Finally, we set *a_η_* = 1,*b_η_* = 1 (E(*η*) = 1/2, Var(*η*) ≃ 1/12) which *a priori* does not push for a sparse graph *G*.

Our model provides us with a rich output that can be summarised in many ways. Regarding the regression structure, one example is that we can use the posterior of the covariate propensity parameter *π_j_, j* = 1,…,*p*, to search for hotspots (i.e., genetic variants that are associated with multiple metabolites). Figure 3 shows the posterior expectations of *π_j_* for each SNP on chromosome 16. In particular, we report rs4985124, rs931406 and most importantly rs12102766 and rs3764261. Analysing the whole binary indicators vector *γ* gives us a lot more information though, as shown in Figure 4, where we plot the marginal posterior inclusion probabilities (mPIP) for each SNP in chromosome 16, all metabolites superimposed. From the plot, we can see how some SNPs are associated with only one or a few outcomes and would thus be missed by only looking at hotspots detection. By thresholding mPIPs at 0.5, we discover a total of 38 associations and the average Bayes FDR (bFDR, see e.g., Lewin et al. (2016)) is ≈ 0.058. The associations found using the mPIP are presented in Supplementary Tables S.3 and S.4. By increasing the mPIP threshold to 0.9, we would keep 32 SNP-metabolite associations with bFDR < 0.01.

**Figure 3:**
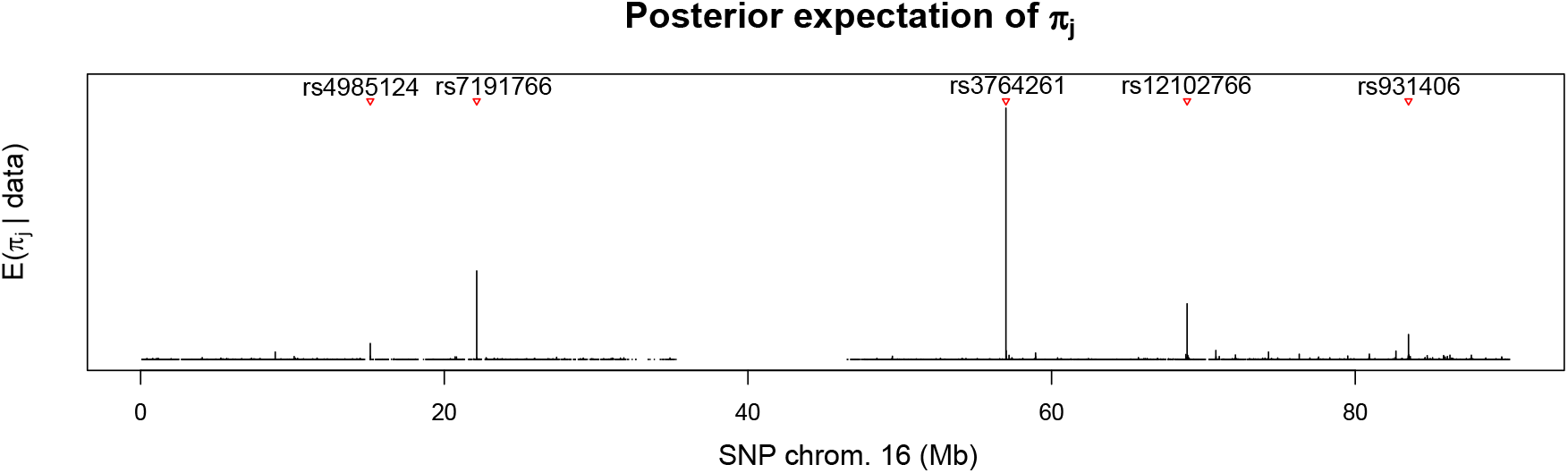
Manhattan plot of the posterior expectation of *π_j, j_* = 1,…, 9310, ordered along chromosome 16, for hotspots detection. Red triangles indicate putative hotspots identified by BayesSUR with the corresponding genetic variant name.

**Figure 4:**
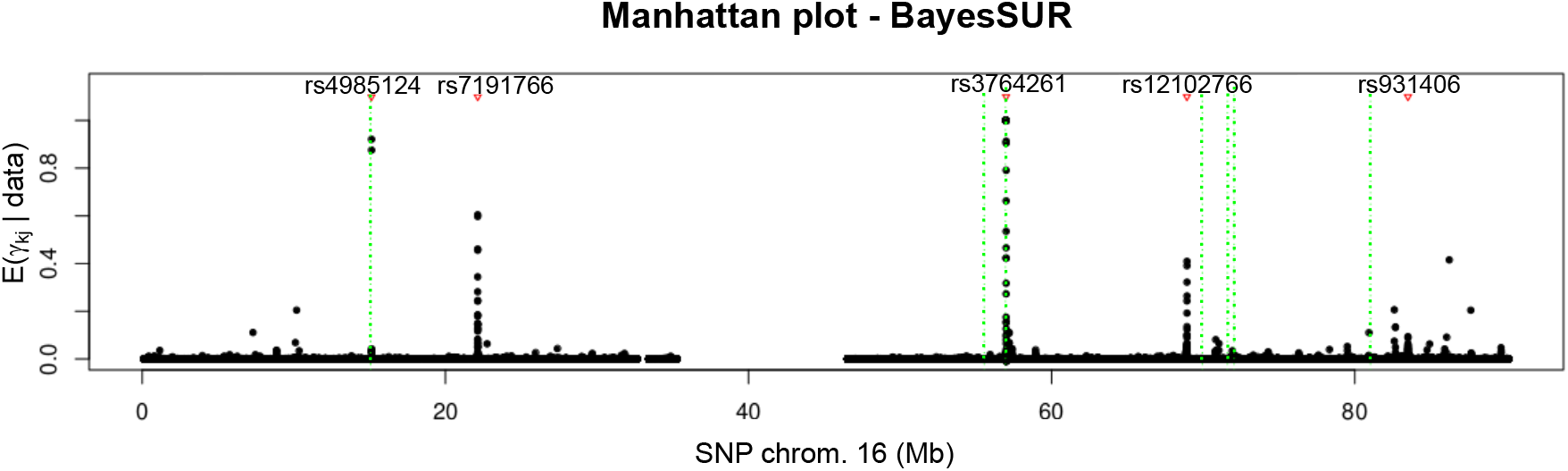
Manhattan plot of marginal posterior inclusion probabilities of association for SNPs in chromosome 16, all metabolites superimposed. Red triangles indicate putative hotspots identified by BayesSUR with the corresponding genetic variant name. Only a few SNPs seem to be relevant and for a restricted number of metabolites. Some SNPs appear to be associated with more than one metabolite. Vertical green dotted lines show previously identified loci associated with lipids metabolites (Kettunen et al., 2016).

The results obtained by BayesSUR confirm the known association between SNP rs3764261 in the *CEPT* gene and HDLs (Sabatti et al., 2009; Kettunen et al., 2012), but additionally highlights the relevance of the *CEPT* locus on different lipoproteins. rs4985124, that we report associated with fatty acids, is situated in the *PDXDC1* locus, roughly 23Kb from SNPs rs11075253 and rs11644601 which were previously linked with fatty acids metabolism (Kettunen et al., 2012, 2016). Finally, rs1210276, that we report associated with multiple metabolites connected to VLDL, is situated in the proximity of rs74249229, previously reported by Kettunen et al. (2016).

A comparison with MatrixeQTL (Shabalin, 2012), a widely used “one-SNP-one-trait-at-a-time” method in GWAS analysis is presented in the Supplementary Material. Not accounting for correlations between metabolites, highly reduces the number of findings in the GWAS analysis. While mostly consistent with BayesSUR in terms of detected loci, multiple close-by SNPs are selected by MatrixeQTL as significantly predictive, whereas BayesSUR method picks only one (see Supplementary Figures S.7 and S.8). Our method, that accounts for residual correlation, was also able to uncover other potential important associations that warrant further investigations, in particular rs7191766 associated with multiple cholesterol-related phenotypes.

Finally, Figure 5 presents a summary of the estimated graph *G*. By thresholding the marginal posterior edge inclusion probabilities (mEPIP) at 0.5, we obtain an adjacency matrix that we represent as a network plot, using the R package `igraph` (Csardi and Nepusz, 2006). In the same plot, we also represent the selected SNPs and their associations with the metabolites.

**Figure 5:**
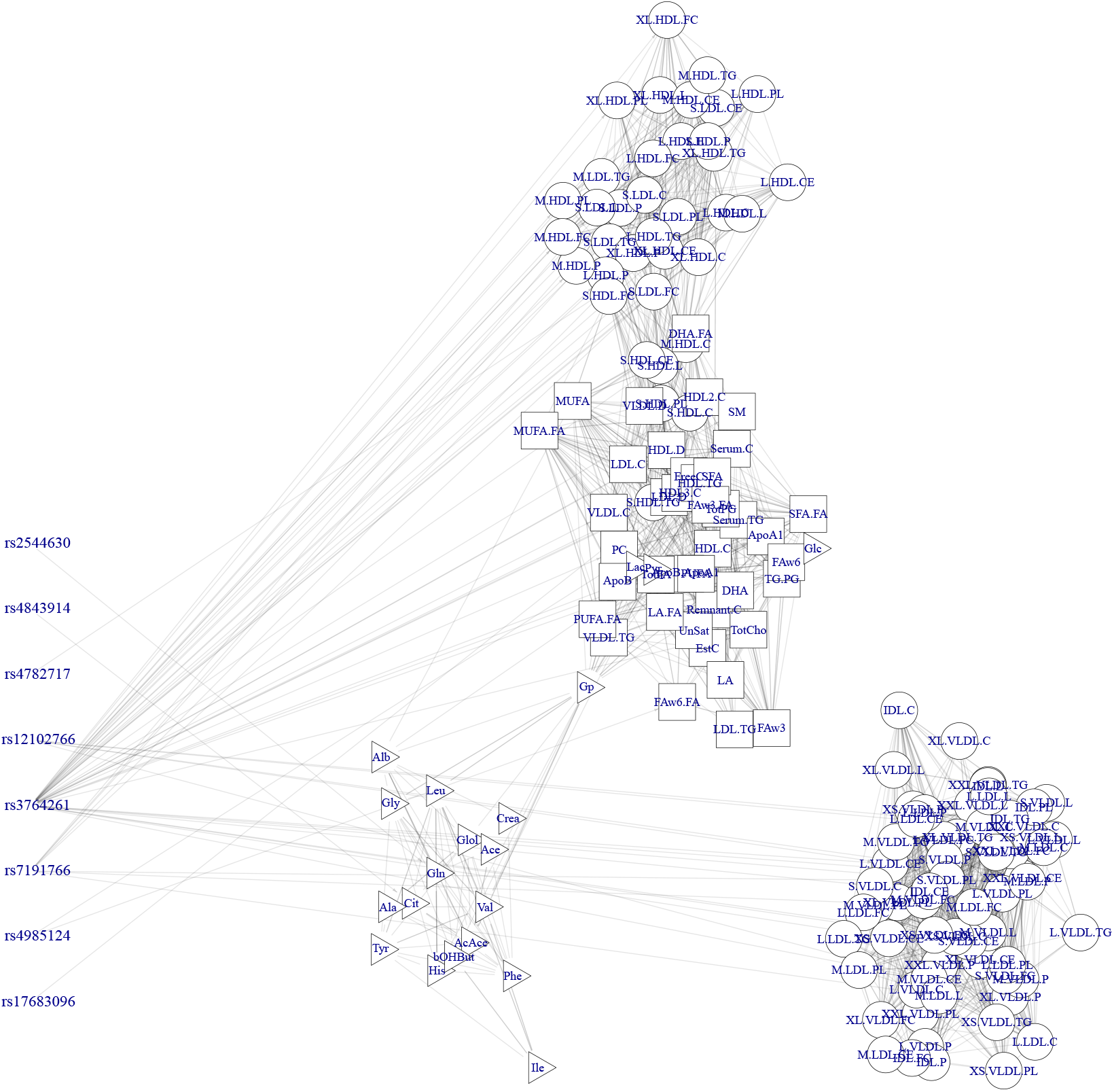
Network representation of the associations between SNPs (right) and metabolite (left) after thresholding the marginal posterior inclusion probabilities at 0.5 and dependence structure between metabolites estimate from the graph *G* after thresholding the marginal posterior edge inclusion probabilities at 0. 5.

An interesting feature of the estimate of *G* is that we recover the three macrogroups mentioned in the Introduction, i.e., lipoprotein concentrations (represented by circles), serum lipids (squares) and low molecular weight metabolites (triangles), with the lipoproteins being further separated into two components. HDLs and LDLs in particular seem to be highly associated with serum lipids, while the VLDL and IDL concentrations form a group almost by themselves. There are associations between the serum lipids and low molecular weight metabolites, driven mostly by a couple of low molecular weight hubs. It is important to note that edges here represent non-zero conditional correlations and we thus expect a much sparser graph than would be seen using marginal correlations. The highly sparse estimate of G also implies that considerable computational gains were achieved using the sparse model.

## 6 Discussion

In this work we present a novel computational method to perform Bayesian variable selection in a multivariate regression setting for Quantitative Trait Loci analysis that takes into account residual correlations between phenotypes while maintaining a flexible association pattern between phenotypes and genotypes. Although conjugacy is lost for this model, by virtue of a crucial reparametrisation of the covariance matrix, our novel results show that: (i) it is possible to obtain simple expressions for the priors distribution of the reparametrised parameters 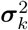 and *ρ_k_, k* = 1,…, *s*, in both dense and sparse cases, (ii) posterior full conditionals are available in closed-form expression, including for the regression coefficients *β_k_*, and (iii) since the likelihood is now computed as a product of independent factors, the posterior updates of *σ_k_*^2^ and *ρ_k_* can be trivially parallelised which greatly increases the computational efficiency of our model. Thus, our method is able to analyse a large number of outcomes and their associations with a large set of predictors, thanks as well to the efficient C++ implementation, as illustrate in the simulation study and in the real data example. It is moreover possible to introduce further computational gains by assuming that the conditional independence structure between the residuals is sparse, and inference on the resulting graph is straightforward to obtain. We demonstrate this feature in a simulated example with 150 responses, where BayesSUR with covariance selection is 30% faster than the version of the algorithm with dense covariance estimation.

We have also shown in the simulation study that, when there is non-negligible residual correlations between the responses, our method exhibits better performance in selecting relevant predictors than existing methods (Bottolo et al., 2011; Lewin et al., 2016) and is able at the same time to effectively perform covariance selection. Computationally, BayeSUR is faster than existing Bayesian sparse SUR methods with covariance selection, although in the simulated examples we haven’t shown the reduction in computational time when multiple cores are used in order to exploit the factorisation of the proposed model. When a large number of responses are considered and the graph is very sparse, BayeSUR computational time is almost comparable to penalized likelihood methods, although the output of the former is much richer (full posterior distributions *versus* point estimates).

Our method is able to scale well in the regime of hundreds of outcomes and thousands of predictors, as demonstrated in the analysis of the NFBC66 mQTL dataset; we are able to recover already published and known associations, as well as uncovering some previously unknown associations that might offer new insights into the relationships between chromosome 16 and lipid metabolism.

One might expect the restriction to decomposable graphs to be too stringent for real applications and various attempt have been made to relax such an assumption, using the *G*-Wishart distribution first introduced by Roverato (2002) (see also Mitsakakis et al. (2011); Wang et al. (2012); Mohammadi and Wit (2015) and references therein for some recent examples). However, the computational disadvantages connected with a general graph are in general exceedingly high (Jones et al., 2005).

The work of Fitch et al. (2014) concludes that, under model assumptions similar to ours, inference on G will asymptotically converge toward minimal triangulations of the true graph, i.e., the decomposable graph with the smallest number of extra edges, and that inference on the covariance matrix is competitive in terms of prediction errors against penalised likelihood methods that estimate unrestricted graphs like the graphical lasso. In practice assuming decomposability seems to be sensible and inference on the covariance matrix under such an assumption sound. Additionally Bayesian modelling averaging enables the estimation via marginal edge inclusion probabilities of a general non-decomposable graph.

Thanks to the very general formulation of the SUR models we expect the present work to find applications beyond the mQTL application presented here, for example in finance, econometrics and other biological settings where linked regression models are already widespread.

## Acknowledgements

The authors are grateful to the associate editor and two anonymous referees for their valuable and detailed comments that greatly improved the presentation of the paper.

## References

Alexopoulos, A. and Bottolo, L. (2020) Bayesian Variable Selection for Gaussian copula regression models. J. Comput. Graph. Stat. In press. 9

Bhadra, A. and Mallick, B. K. (2013) Joint high-dimensional Bayesian variable and covariance selection with an application to eQTL analysis. Biometrics, 69, 447–457. 2, 5, 11

Bottolo, L., Petretto, E., Blankenberg, S., Cambien, F., Cook, S. A., Tiret, L. and Richardson, S. (2011) Bayesian detection of expression quantitative trait loci hot-spots. Genetics, 189, 1449–1459. 3, 5, 9, 11, 12, 17, 20

Bottolo, L. and Richardson, S. (2010) Evolutionary stochastic search for Bayesian model exploration. Bayesian Anal., 5, 583–618. 9

Brown, P. J., Vannucci, M. and Fearn, T. (1998) Multivariate Bayesian variable selection and prediction. J. R. Stat. Soc. Series B Stat. Methodol., 60, 627–641. 2 —

Brown, P. J., Vannucci, M. and Fearn, T.(2002)Bayes model averaging with selection of regressors. J. R. Stat. Soc. Series B Stat. Methodol., 64, 519–536. 2

Carvalho, C. M., Massam, H. and West, M. (2007) Simulation of hyper-inverse Wishart distributions in graphical models. Biometrika, 94, 647–659. 6, 7

Chib, S. (1995) Marginal likelihood from the Gibbs output. J. Am. Stat. Assoc., 90, 1313–1321. 3, 13

Cichonska, A., Rousu, J., Marttinen, P., Kangas, A. J., Soininen, P., Lehtimäki, T., Raitakari, O. T., Järvelin, M.-R., Salomaa, V., Ala-Korpela, M. et al. (2016) metaCCA: Summary statistics-based multivariate meta-analysis of genome-wide association studies using canonical correlation analysis. Bioinformatics, 32, 1981–1989. 3

Csardi, G. and Nepusz, T. (2006) The igraph software package for complex network research. Inter-Journal - Complex Systems, 1695. 18

Datta, A., Banerjee, S., Hodges, J. S. and Gao, L. (2019) Spatial disease mapping using Directed Acyclic Graph Auto-Regressive (DAGAR) models. Bayesian Anal., 14, 1221–1244. 3

Dawid, A. P. (1981) Some matrix-variate distribution theory: notational considerations and a Bayesian application. Biometrika, 68, 265–274. 6

Dempster, A. P. (1969) Elements of Continuous Multivariate Analysis. Addison Wesley Longman, Boston. 6

Fitch, A. M., Jones, M. B. and Massam, H. (2014) The performance of covariance selection methods that consider decomposable models only. Bayesian Anal., 9, 659–684. 12, 20

Fusi, N., Stegle, O. and Lawrence, N. D. (2012) Joint modelling of confounding factors and prominent genetic regulators provides increased accuracy in genetical genomics studies. PLoS Comput. Biol., 8, e1002330. 2

George, E. and McCulloch, R. E. (1993) Variable selection via Gibbs sampling. J. Am. Stat. Assoc., 88, 881–889. 3, 4, 13

Giri, N. C. (2014) Multivariate Statistical Inference. Academic Press, London. 5

Giudici, P. and Green, P. J. (1999) Decomposable graphical Gaussian model determination. Biometrika, 86, 785–801. 16

Green, P. and Thomas, A. (2013) Sampling decomposable graphs using a Markov chain on junction trees. Biometrika, 100, 91–110. 3, 7, 10, 16

Holmes, C. C., Denison, D. G. T. and Mallick, B. K. (2002) Accounting for model uncertainty in seemingly unrelated regressions. J. Comput. Graph. Stat., 11, 533–551. 5, 8

Holmes, C. C. and Held, L. (2006) Bayesian auxiliary variable models for binary and multinomial regression. Bayesian Anal., 1, 145–168. 9

Inouye, M., Ripatti, S., Kettunen, J., Lyytikäinen, L.-P., Oksala, N., Laurila, P.-P., Kangas, A. J., Soininen, P., Savolainen, M. J., Viikari, J. et al. (2012) Novel loci for metabolic networks and multi-tissue expression studies reveal genes for atherosclerosis. PLoS Genet., 8, e1002907. 2

Jia, Z. and Xu, S. (2007) Mapping quantitative trait loci for expression abundance. Genetics, 176, 611–623. 3

Jones, B., Carvalho, C., Dobra, A., Hans, C., Carter, C. and West, M. (2005) Experiments in stochastic computation for high-dimensional graphical models. Stat. Sci., 20, 388–400. 20

Kettunen, J., Demirkan, A., Würtz, P., Draisma, H. H., Haller, T., Rawal, R., Vaarhorst, A., Kangas, A. J., Lyytikäinen, L.-P., Pirinen, M. et al. (2016) Genome-wide study for circulating metabolites identifies 62 loci and reveals novel systemic effects of LPA. Nat. Commun., 7, 11122. 16, 17, 18

Kettunen, J., Tukiainen, T., Sarin, A.-P., Ortega-Alonso, A., Tikkanen, E., Lyytikäinen, L.-P., Kangas, A. J., Soininen, P., Würtz, P., Silander, K. et al. (2012) Genome-wide association study identifies multiple loci influencing human serum metabolite levels. Nature Genet., 44, 269. 2, 17

Lauritzen, S. L. (1996) Graphical Models. Oxford University Press, New York. 6

Lewin, A., Saadi, H., Peters, J. E., Moreno-Moral, A., Lee, J. C., Smith, K. G. C., Petretto, E., Bottolo, L. and Richardson, S. (2016) MT-HESS: An efficient Bayesian approach for simultaneous association detection in OMICS datasets, with application to eQTL mapping in multiple tissues. Bioinformatics, 32, 523–532. 9, 12, 17, 20

Liang, F. and Wong, W. H. (2000) Evolutionary Monte Carlo: Applications to *C_p_* model sampling and change point problem. Stat. Sin., 10, 317–342. 9

Marttinen, P., Pirinen, M., Sarin, A.-P., Gillberg, J., Kettunen, J., Surakka, I., Kangas, A. J., Soininen, P., O’Reilly, P., Kaakinen, M. et al. (2014) Assessing multivariate gene-metabolome associations with rare variants using Bayesian reduced rank regression. Bioinformatics, 30, 2026–2034. 2

Mitsakakis, N., Massam, H. and Escobar, M. D. (2011) A Metropolis-Hastings based method for sampling from the G-Wishart distribution in Gaussian graphical models. Electron. J. Stat., 5, 18–30. 20

Mohammadi, A. and Wit, E. C. (2015) Bayesian structure learning in sparse Gaussian graphical models. Bayesian Anal., 10, 109–138. 20

Mohammadi, A. and Wit, E. C. (2019) BDgraph: An R package for Bayesian structure learning in graphical models. J. Stat. Softw., 89, 1–30. 12

Paulsen, V. I., Power, S. C. and Smith, R. R. (1989) Schur products and matrix completions. J. Funct. Anal., 85, 151–178. 6

Petretto, E., Bottolo, L., Langley, S. R., Heinig, M., McDermott-Roe, C., Sarwar, R., Pravenec, M., Hübner, N., Aitman, T. J., Cook, S. A. et al. (2010) New insights into the genetic control of gene expression using a Bayesian multi-tissue approach. PLoS Comput. Biol., 6, e1000737. 2

Pourahmadi, M. (1999) Joint mean-covariance models with applications to longitudinal data: Unconstrained parameterisation. Biometrika, 86, 677–690. 3

Richardson, S., Bottolo, L. and Rosenthal, J. S. (2010) Bayesian models for sparse regression analysis of high dimensional data. In Bayesian Statistics (eds. J. M. Bernardo, M. J. Bayarri, J. O. Berger, A. P. Dawid, D. Heckerman, A. F. M. Smith and M. West), vol. 9, 539–569. Oxford University Press, New York. 11

Roberts, G. O. and Rosenthal, J. S. (2009) Examples of adaptive MCMC. J. Comput. Graph. Stat., 18, 349–367. 10

Rodriguez-Martinez, A., Posma, J. M., Ayala, R., Neves, A. L., Anwar, M., Petretto, E., Emanueli, C., Gauguier, D., Nicholson, J. K. and Dumas, M.-E. (2017) MWASTools: an R/Bioconductor package for metabolome-wide association studies. Bioinformatics, 34, 890–892. 3

Rothman, A. J., Levina, E. and Zhu, J. (2010) Sparse multivariate regression with covariance estimation. J. Comput. Graph. Stat., 19, 947–962. 4, 14, 15

Roverato, A. (2000) Cholesky decomposition of a hyper inverse Wishart matrix. Biometrika, 87, 99–112. 6

Roverato, A. (2002) Hyper inverse Wishart distribution for non-decomposable graphs and its application to Bayesian inference for Gaussian graphical models. Scand. J. Stat., 29, 391–411. 20

Ruffieux, H., Davison, A. C., Hager, J., Inshaw, J., Fairfax, B. P., Richardson, S. and Bottolo, L. (2020a) A global-local approach for detecting hotspots in multiple-response regression. Ann. Appl. Stat., 14, 905–928. 3

Ruffieux, H., Davison, A. C., Hager, J. and Irincheeva, I. (2017) Efficient inference for genetic association studies with multiple outcomes. Biostatistics, 18, 618–636. 3

Ruffieux, H., Fairfax, B. P., Nassiri, I., Vigorito, E., Wallace, C., Richardson, S. and Bottolo, L. (2020b) EPISPOT: An epigenome-driven approach for detecting and interpreting hotspots in molecular QTL studies. bioRxiv. URL: https://doi.org/10.1101/2020.09.21.305789. 3

Sabatti, C., Service, S. K., Hartikainen, A.-L., Pouta, A., Ripatti, S., Brodsky, J., Jones, C. G., Zaitlen, N. A., Varilo, T., Kaakinen, M. et al. (2009) Genome-wide association analysis of metabolic traits in a birth cohort from a founder population. Nature Genet., 41, 35. 17

Scott-Boyer, M. P., Imholte, G. C., Tayeb, A., Labbe, A., Deschepper, C. F. and Gottardo, R. (2012) An integrated hierarchical Bayesian model for multivariate eQTL mapping. Stat. Appl. Genet. Mol. Biol., 11. 3

Shabalin, A. (2012) Matrix eQTL: Ultra fast eQTL analysis via large matrix operations. Bioinformatics, 28, 1353–1358. 17

Soininen, P., Kangas, A. J., Würtz, P., Tukiainen, T., Tynkkynen, T., Laatikainen, R., Järvelin, M.-R., Kähönen, M., Lehtimäki, T., Viikari, J. et al. (2009) High-throughput serum NMR metabonomics for cost-effective holistic studies on systemic metabolism. Analyst, 134, 1781–1785. 2

Stingo, F. and Marchetti, G. (2014) Efficient local updates for undirected graphical models. Stat. Comput., 25, 159–171. 3

Thomas, A. and Green, P. J. (2009) Enumerating the junction trees of a decomposable graph. J. Comput. Graph. Stat., 18, 930–940. 7

Wang, H. (2010) Sparse seemingly unrelated regression modelling: Applications in finance and econometrics. Comput. Stat. Data Anal., 54, 2866–2877. 3, 4, 13, 16

Wang, H. et al. (2012) Bayesian graphical lasso models and efficient posterior computation. Bayesian Anal., 7, 867–886. 3, 20

Wermuth, N. (1980) Linear recursive equations, covariance selection, and path analysis. J. Am. Stat. Assoc., 75, 963–972. 3

Würtz, P., Kangas, A. J., Soininen, P., Lawlor, D. A., Davey Smith, G. and Ala-Korpela, M. (2017) Quantitative Serum Nuclear Magnetic Resonance Metabolomics in Large-Scale Epidemiology: A Primer on-Omic Technologies. American Journal of Epidemiology, 186, 1084–1096. URL: https://doi.org/10.1093/aje/kwx016. 16

Zellner, A. and Ando, T. (2010) A direct Monte Carlo approach for Bayesian analysis of the seemingly unrelated regression model. J. Econom., 159, 33–45. 3, 9

