## Supplementary Material for "A computationally efficient Bayesian Seemingly Unrelated Regressions model for high-dimensional Quantitative Trait Loci discovery"

### S.1 Technical notes

#### S.1.1 Reparametrisation and prior derivation

In this section we use decomposable graphs to model sparsity and show that the new parameters $\boldsymbol{\rho}_k$, $k = 1, \ldots, s$, are simply related to the graph structure and, hence, retain useful computational properties for MCMC estimation of the Seemingly Unrelated Regressions (SUR) models.

##### Perfect elimination order and notation

We start by introducing some useful graph concepts, alongside the notation that we will use in the rest of the Supplementary Material. A separator of a graph $G$ is a completely-connected subgraph of $G$ which separates the graph into two components such that any path between the two components must pass through the separator. The two components and the separator form a decomposition of $G$. The set of subgraphs which cannot be further decomposed form the prime components of the graph. If all the prime components are complete then the graph is said to be decomposable. In this case the prime components $P_q$ can be ordered in such a way that for every $q > 1$ there exists $m < q$ such that $P_q \cap H_q \subset P_m$, where $H_q = \bigcup_{l=1}^{q-1} P_l$ for $q = 2, \ldots, Q$. This is called the "running intersection property" (Lauritzen, 1996). With this notation, we also define the separators $S_q = P_q \cap H_q$ and residuals $R_q = P_q \setminus S_q$ for $q = 2, \ldots, Q$. The separators $\{S_2, \ldots, S_Q\}$ are not necessarily distinct from each other.

The (non-unique) ordering of the prime components is called a "perfect ordering" and has the property that nodes in residual $R_q$ are independent of nodes in $H_q \setminus S_q$, conditionally on the rest of the graph and therefore the elements of the precision matrix relating pairs of nodes in those sets are structurally equal to zero. The so-called "perfect elimination ordering" of nodes (also non-unique, even for a given perfect ordering of prime components) will be denoted $\xi$. This is defined by taking the nodes in $P_1$ in any order, followed by the nodes in $R_2$ in any order and so on for the rest of the residuals up to $R_Q$.

Figure S.1 shows a simple example of a small decomposable graph. One possible perfect

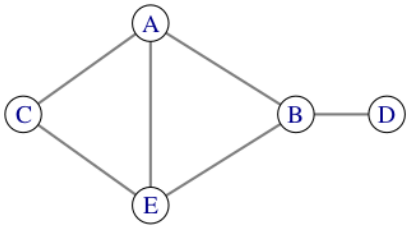


Figure S.1: A toy decomposable graph, used to exemplify perfect ordering of its cliques and the perfect elimination ordering.

ordering of the prime components is $P_1 = \{A, B, E\}$, $P_2 = \{A, C, E\}$, $P_3 = \{B, D\}$, with separators $S_2 = \{A, E\}$ and $S_3 = \{B\}$. Given this ordering of the prime components, a possible perfect elimination ordering of the nodes is $\{A, B, E, C, D\}$.

With the variables ordered according to the perfect elimination ordering, we have that $\Lambda_{kl} = 0 \implies \rho_{kl} = 0$, $k, l = 1, \ldots, s$, $k \neq l$, where $\Lambda$ is the precision matrix $C^{-1}$. This was proved by Paulsen et al. (1989) (a short proof in our notation is also given in the Supplementary Material Section S.1.3). This means that the reparametrisation of the covariance matrix is a convenient way to encode the conditional independence between variables. In addition, we see that conditionally on a given sparse graph, we will not need to estimate values of $\rho_{kl}$ for many pairs of variables, allowing for computational savings. More specifically, given a particular decomposable graph structure, $\rho_{kl}$ are only estimated for pairs of variables $k, l$ contained in the same prime component of the graph.

##### Reparametrisation definition

Here, we show further that the new parameters $\sigma_k^2$ and $\boldsymbol{\rho}_k$, $k = 1, \ldots, s$, are defined straightforwardly in terms of the covariance matrices of the prime components of the graph and, hence, we calculate priors in the reparametrised space. Assuming the perfect elimination ordering, take any prime component $P_q$ and define variable $k$ to be the final variable in the residual $R_q$. With this ordering of variables, $\boldsymbol{\rho}_k$ relates variable $k$ to the set of all variables in prime components $1, \ldots, q$, which is the set of nodes in the subgraph $H_{q+1}$. Thus, from the definition of the reparametrisation, $\boldsymbol{\rho}_k = -\boldsymbol{\omega}_k \omega_k^{-1}$, where $\boldsymbol{\omega}_k$ is the final column of $C_{H_{q+1}}^{-1}$ without the last element and $\omega_k$ its last diagonal entry.

The marginal precision matrix for the subgraph $H_{q+1}$, consisting of all prime components from 1 to $q$, can be written

$$C_{H_{q+1}}^{-1} = \begin{pmatrix} \Omega_{H_q\setminus S_q} & \Omega_{H_q\setminus S_q,S_q} & 0 \\ \Omega_{H_q\setminus S_q,S_q}^T & \Omega_{S_q} & \Omega_{S_q,R_q} \\ 0 & \Omega_{S_q,R_q}^T & \Omega_{R_q} \end{pmatrix} \tag{S.1}$$

as illustrated later, which shows that the elements of $\boldsymbol{\rho}_k$ corresponding to variables in $H_q\setminus S_q$ are zero. Further block matrix calculation shows that

$$C_{P_q}^{-1} = \begin{pmatrix} \Omega_{S_q} - \Omega_{H_q,S_q}^T \Omega_{H_q\setminus S_q}^{-1} \Omega_{H_q,S_q} & \Omega_{S_q,R_q} \\ \Omega_{S_q,R_q}^T & \Omega_{R_q} \end{pmatrix}, \tag{S.2}$$

where we highlight that the off-diagonal blocks in $C_{P_q}^{-1}$ appear unmodified also in $C_{H_{q+1}}^{-1}$ thanks to the zeros in the latter.

Now, let's define $\boldsymbol{\rho}_{k,P_q}$ for the non-zero elements of $\boldsymbol{\rho}_k$. From Equations (S.1) and (S.2), we can see that this can be written in terms of the final column of $C_{P_q}^{-1}$ without the last element and the last diagonal entry of $C_{P_q}^{-1}$. Hence, again by block matrix manipulation, we can write

$$\boldsymbol{\rho}_{k,P_q} = (C_{P_q}^{(k-1)})^{-1} \boldsymbol{c}_{k,P_q},$$

where $C_{P_q}^{(k-1)}$ is the sub-matrix of $C_{P_q}$ with variable $k$ removed and $\boldsymbol{c}_{k,P_q}$ is the final column of $C_{P_q}$ without the last element. This is the same as the original definition of $\boldsymbol{\rho}_k$, except that, due to the structural zeros in the precision matrix, now we only need to use the sub-matrix $C_{P_q}$ corresponding to the prime component $P_q$ containing $R_q$.

Then, consider

$$\begin{aligned} \sigma_k^2 &= c_k - \boldsymbol{c}_k^T C_{(k-1)}^{-1} \boldsymbol{c}_k = c_k - \boldsymbol{\rho}_k^T C_{(k-1)} \boldsymbol{\rho}_k \\ &= c_k - \boldsymbol{\rho}_{k,P_q}^T C_{P_q}^{(k-1)} \boldsymbol{\rho}_{k,P_q} = c_k - \boldsymbol{c}_{k,P_q}^T (C_{P_q}^{(k-1)})^{-1} \boldsymbol{c}_{k,P_q}. \end{aligned}$$

This shows that $\sigma_k^2$ also has the same definition in terms of the sub-covariance matrix $C_{P_q}$.

Thus, we see that with a perfect elimination ordering of variables, the new parameters $\sigma_k^2$ and $\boldsymbol{\rho}_k$, are completely specified by the marginal covariance matrices of the prime components of the decomposable graph.

Due to the conditional independence structures, the distribution of the entire covariance matrix can be decomposed over the prime components and separators (Carvalho et al., 2007):

$$\begin{aligned} p(C) &= p(C_{P_1}) \prod_{q=2}^{Q} p(C_{P_q} \mid C_{S_q}) \delta(C_{H_q \setminus S_q, R_q} = \cdot) \\ &= p(C_{P_1}) \prod_{q=2}^{Q} p(C_{R_q}, C_{S_q, R_q} \mid C_{S_q}) \delta(C_{H_q \setminus S_q, R_q} = \cdot), \end{aligned}$$

where $C_{P_q}$ and $C_{S_q}$ stand for the covariance matrices of $P_q$ and $S_q$, respectively, $C_{S_q,R_q}$ is the cross-covariance between the separator and residual of component $q$ and $C_{H_q \setminus S_q, R_q}$ is the cross-covariance between residual $R_q$ and the part of the graph lower in the ordering than component $q$ and separated from it: $H_q \setminus S_q$. The $\delta(C_{H_q \setminus S_q, R_q} = \cdot)$ is the completion operator (Carvalho et al., 2007), which uses the structural zeros in the precision matrix defined by the graph to fill in the remaining elements of the covariance matrix.

Carvalho et al. (2007) shows that this can be reparametrised in terms of the Schur complements of the covariance matrices of the residuals $C_{R_q}$ and the cross-covariances between the separators and residuals $C_{S_q,R_q}$, with these quantities having Inverse-Wishart and Normal densities. Our new parametrisation has a similar structure, since the $\sigma_k^2$ are univariate Schur complements of sub-matrices of the covariance matrix and the $\boldsymbol{\rho}_k$ are related to the cross-covariances of the same sub-matrices.

##### Priors in the reparametrised space

For decomposable graphs, a convenient prior distribution for the covariance matrix is the Hyper Inverse-Wishart (HIW), which is the conjugate prior for each prime component in a Gaussian graphical model. The Hyper Inverse-Wishart $C \sim \text{HIW}_G(\nu, M)$ is defined as the distribution such that the covariance matrix for each prime component is marginally

Inverse-Wishart, i.e., $C_{P_q} \sim \mathrm{IW}(\nu - (s - |P_q|), M_{P_q})$, where $M_{P_q}$ is the sub-matrix of $M$ corresponding to $P_q$ and $|P_q|$ is its cardinality.

For clarity, we relabel variables conditional on a given decomposable graph structure. To obtain the perfect elimination ordering of variables discussed above, we label variables with index $t = (|S_q|{+}1), \ldots, |P_q|$ within each residual $R_q$ for components $q = 2, \ldots, Q$. For the first prime component, all variables are labelled $t = 1, \ldots, |P_1|$. As mentioned in Section 2 of the main text, with this labelling, the HIW prior on the covariance matrix is transformed to

$$p(\sigma_{11}^2) \prod_{t=2}^{|P_1|} p(\sigma_{1t}^2) p(\boldsymbol{\rho}_{1t} \mid \sigma_{1t}^2) \prod_{q=2}^{Q} \prod_{t=1}^{|R_q|} p(\sigma_{qt}^2) p(\boldsymbol{\rho}_{qt} \mid \sigma_{qt}^2),$$

where the densities are

$$\begin{aligned} \sigma_{qt}^2 &\sim \mathrm{IGa}((\nu - s + t + |S_q|)/2, (m_{qt} - \boldsymbol{m}_{qt}^T M_{P_q,(t-1)}^{-1} \boldsymbol{m}_{qt})/2), \\ \boldsymbol{\rho}_{qt} | \sigma_{qt}^2 &\sim \mathrm{N}(M_{P_q,(t-1)}^{-1} \boldsymbol{m}_{qt}, \sigma_{qt}^2 M_{P_q,(t-1)}^{-1}) \end{aligned}$$

for all $q$ and $t$ except $\sigma_{11}^2 \sim \mathrm{IGa}((\nu - s + 1)/2, m_{11}/2)$.

An important feature of this transformation is that we do not need to do the completion operation to fill in the covariances between the separated parts of prime components.

##### S.1.2 Reparametrisation in terms of the precision matrix

Here, we derive the reparametrisation in terms of the precision matrix $\Lambda = C^{-1}$. This will be usefull to show that zeros in the precision matrix, conditioned on a decomposable graph, directly translate to zeros in the coefficients $\boldsymbol{\rho}$. We can decompose the precision matrix $\Lambda$ in the same way as we decompose $C$, i.e.,

$$\Lambda_{(k)} = \begin{pmatrix} \Lambda_{(k-1)} & \boldsymbol{\lambda}_k \\ \boldsymbol{\lambda}_k^T & \lambda_k \end{pmatrix}$$

for all $k = 2, \ldots, s$, with $\Lambda_{(s)} = \Lambda$ and $\Lambda_{(1)} = \lambda_1$ and $\boldsymbol{\lambda}_1$ null. Note that $\Lambda_{(k)}$ is not the marginal precision matrix for responses $1, \ldots, k$. For this we use instead the different decomposition

$$C_{(k)}^{-1} = \begin{pmatrix} \Omega_{(k-1)} & \boldsymbol{\omega}_k \\ \boldsymbol{\omega}_k^T & \omega_k \end{pmatrix} \tag{S.3}$$

for all $k = 2, \ldots, s$, with $\Omega_{(s)} = \Lambda$, $\Omega_{(s-1)} = \Lambda_{(s-1)}$, $\Omega_{(1)} = \omega_1$ and $\boldsymbol{\omega}_1$ is null.

Recall the original decomposition in Section 2.1 of the main text

$$C_{(k)} = \begin{pmatrix} C_{(k-1)} & \boldsymbol{c}_k \\ \boldsymbol{c}_k^T & c_k \end{pmatrix}. \tag{S.4}$$

Using standard block matrix inverse identities, we see that the decompositions in Equations (S.3) and (S.4) are related by

$$\begin{aligned} C_{(k-1)}^{-1}\boldsymbol{c}_k &= -\boldsymbol{\omega}_k \omega_k^{-1} \\ c_k - \boldsymbol{c}_k^T C_{(k-1)}^{-1}\boldsymbol{c}_k &= \omega_k^{-1}. \end{aligned}$$

Hence, we can write down the new parameters in the transformed space as

$$\begin{aligned} \sigma_k^2 &= \omega_k^{-1}, \quad k = 1, \ldots, s, \\ \boldsymbol{\rho}_k &= -\boldsymbol{\omega}_k \omega_k^{-1}, \quad k = 2, \ldots, s, \end{aligned}$$

with $\boldsymbol{\omega}_s = \boldsymbol{\lambda}_s$ and $\omega_s = \lambda_s$. So we notice that the new parameters are directly related to the marginal precision matrices of the subsets of response variables and hence the $\boldsymbol{\rho}$ directly relate to the partial correlations amongst the subsets of variables with the remaining variables marginalised out.

We can use the Cholesky decomposition of $C$ to derive a relation between the new parameters and the complete precision matrix $\Lambda$. Let $P$ be a lower triangular matrix with $P_{kl} = \rho_{kl}$ $(k > l)$ and zeros on the diagonal. The reparametrisation of the covariance matrix converts

$$\boldsymbol{y}_k = X_{\boldsymbol{\gamma}_k}\boldsymbol{\beta}_{\boldsymbol{\gamma}_k} + \boldsymbol{u}_k, \quad \boldsymbol{u}_k \sim \mathrm{N}\left(\mathbf{0}, C\right),$$

into

$$\boldsymbol{y}_k = X_{\boldsymbol{\gamma}_k}\boldsymbol{\beta}_{\boldsymbol{\gamma}_k} + P\boldsymbol{u}_k + \boldsymbol{\varepsilon}_k, \quad \boldsymbol{\varepsilon}_k \sim \mathrm{N}\left(\mathbf{0}, S\right),$$

where $S$ is a diagonal matrix such that $S_{kk} = \sigma_k^2$. Thus, we see that

$$\boldsymbol{u}_k = \left(\mathbb{I}_n - P\right)^{-1}\boldsymbol{\varepsilon}_k \sim \mathrm{N}\left(\mathbf{0}, \left(\mathbb{I}_n - P\right)^{-1} S\{(\mathbb{I}_n - P)^{-1}\}^T\right).$$

Given that $C$ is symmetric and positive definite, there exists a unique modified Cholesky decomposition such that $C = LDL^T$. We thus observe that the reparametrisation is related to the Cholesky decomposition with $L = \left(\mathbb{I}_n - P\right)^{-1}$ and $D = S$. Then, we can write the precision matrix in terms of the new parameters

$$\Lambda = C^{-1} = (\mathbb{I} - P)^T S^{-1}(\mathbb{I} - P)$$

from which we can see that

$$\boldsymbol{\lambda}_k = -\sigma_k^{-2}\boldsymbol{\rho}_k + \sum_{m=k+1}^{s} \sigma_m^{-2}\rho_{m,k}\boldsymbol{\rho}_{m[1:k-1]}$$

for $k = 2, \ldots, s$.

##### S.1.3 Proof that zeros in $\boldsymbol{\rho}_k$ correspond to zeros in $\Lambda$

The decomposable graph structure means that the precision matrix $\Lambda$ for the entire graph can be written as

$$\Lambda = \begin{pmatrix} \Lambda_{H_Q \setminus S_Q} & \Lambda_{H_Q \setminus S_Q, S_Q} & 0 \\ \Lambda^T_{H_Q \setminus S_Q, S_Q} & \Lambda_{S_Q} & \Lambda_{S_Q, R_Q} \\ 0 & \Lambda^T_{S_Q, R_Q} & \Lambda_{R_Q} \end{pmatrix},$$

where $Q$ is the final prime component in the ordering above. The blocks of zeros are due to the conditional independence defined by the graph structure, since by definition $R_Q$ is conditionally independent of $H_Q \setminus S_Q$ given the separator $S_Q$.

With the perfect elimination ordering, variable $s$ is the final variable in the residual of the final prime component $R_Q$, corresponding to the final row/column in $\Lambda$ as ordered here. For the variable $s$, we know from standard matrix operations that

$$\boldsymbol{\rho}_s = -\boldsymbol{\lambda}_s \lambda_s^{-1},$$

so we can see here that all elements of the vector $\boldsymbol{\rho}_s$ corresponding to the partial correlations between variable $s$ and variables in $H_Q \setminus S_Q$ are zero. Next, consider the remaining variables in $R_Q$, labelling these $s-1, s-2, \ldots$. For any variable $k$, we have the recursion formula relating $k$ to variables $m > k$

$$\sigma_k^{-2} \boldsymbol{\rho}_k = \sum_{m>k} \sigma_m^{-2} \rho_{mk} \boldsymbol{\rho}_m - \boldsymbol{\lambda}_k.$$

From this, it is easy to see that for any variable $k$ in the final residual $R_Q$, the zeros in $\boldsymbol{\rho}_k$ are directly inherited from the $\boldsymbol{\rho}_k, \boldsymbol{\rho}_{k+1}, \ldots, \boldsymbol{\rho}_s$. All these variables have the same zeros in the $\boldsymbol{\lambda}_k, \boldsymbol{\lambda}_{k+1}, \ldots, \boldsymbol{\lambda}_s$ sub-vectors of the precision matrix, so for all variables in $R_Q$, the vector $\boldsymbol{\rho}_k$ has zeros where $\boldsymbol{\lambda}_k$ has zeros, corresponding to partial correlations with variables in $H_Q \setminus S_Q$.

Now, let's consider the residual of the next to final prime component, $R_{Q-1}$. By definition of the decomposable graph structure, the variables in $R_{Q-1}$ are separated from variables in $H_{Q-1} \setminus S_{Q-1}$ and so $\Lambda_{H_{Q-1}\setminus S_{Q-1}, R_{Q-1}} = 0$. The running intersection property of a decomposable graph implies that $R_Q$ must be separated from at least one of $R_{Q-1}$ and $H_{Q-1} \setminus S_{Q-1}$. If $R_Q$ is separated from $R_{Q-1}$ then $\rho_{mk} = 0$ for all $m \in R_Q$ and $k \in R_{Q-1}$. If $R_Q$ is separated from $H_{Q-1} \setminus S_{Q-1}$ then $\lambda_{kl} = 0 \implies \lambda_{ml} = 0 \implies \rho_{ml} = 0$ for all $m \in R_Q$ and $k \in R_{Q-1}$. In either case this leads to the conclusion that $\lambda_{kl} = 0 \implies \rho_{kl} = 0$. Hence, again, $\boldsymbol{\rho}_k$ has zeros where $\boldsymbol{\lambda}_k$ has zeros, corresponding to partial correlations between variables in $R_{Q-1}$ and variables in $H_{Q-1} \setminus S_{Q-1}$.

The same argument applies to all successive residuals of the graph, so we see that for all variables, $\rho_{kl} = 0$ where $\Lambda_{kl} = 0$.

##### S.1.4 Marginal precision matrix for subgraph

Here, we consider decomposable graphs. We show that the marginal precision matrix $C_{H_q}^{-1}$ for each subgraph $H_q$ has the same zero structure as the corresponding block of the precision matrix for the entire graph $\Lambda_{H_q}$. First, we can partition the precision matrix into the final residual and the rest of the graph:

$$\Lambda = \begin{pmatrix} \Lambda_{H_Q} & \Lambda_{H_Q,R_Q} \\ \Lambda_{H_Q,R_Q}^T & \Lambda_{R_Q} \end{pmatrix}$$

conformably with the covariance matrix

$$C = \begin{pmatrix} C_{H_Q} & C_{H_Q,R_Q} \\ C_{H_Q,R_Q}^T & C_{R_Q} \end{pmatrix}$$

and from these we have

$$C_{H_Q}^{-1} = \Lambda_{H_Q} - \Lambda_{H_Q,R_Q} \Lambda_{R_Q}^{-1} \Lambda_{H_Q,R_Q}^T. \tag{S.5}$$

Next, we partition $\Lambda_{H_Q}$ into the next residual $R_{Q-1}$, its separator $S_{Q-1}$ and the remainder

$$\Lambda_{H_Q} = \begin{pmatrix} \Lambda_{H_{Q-1}\setminus S_{Q-1}} & \Lambda_{H_{Q-1}\setminus S_{Q-1},S_{Q-1}} & 0 \\ \Lambda^T_{H_{Q-1}\setminus S_{Q-1},S_{Q-1}} & \Lambda_{S_{Q-1}} & \Lambda_{S_{Q-1},R_{Q-1}} \\ 0 & \Lambda^T_{S_{Q-1},R_{Q-1}} & \Lambda_{R_{Q-1}} \end{pmatrix},$$

where the zero blocks appear by definition of the decomposability of the graph.

Now, let's calculate the second term in Equation (S.5)

$$\Lambda_{H_Q,R_Q}\Lambda^{-1}_{R_Q}\Lambda^T_{H_Q,R_Q} = \begin{pmatrix} \Lambda_{H_{Q-1}\setminus S_{Q-1},R_Q} \\ \Lambda_{S_{Q-1},R_Q} \\ \Lambda_{R_{Q-1},R_Q} \end{pmatrix} \Lambda^{-1}_{R_Q} \begin{pmatrix} \Lambda_{H_{Q-1}\setminus S_{Q-1},R_Q} & \Lambda_{S_{Q-1},R_Q} & \Lambda_{R_{Q-1},R_Q} \end{pmatrix}.$$

The running intersection property of a decomposable graph implies that either (a) $R_Q$ is separated from $R_{Q-1}$ or (b) $R_Q$ is separated from $H_{Q-1} \setminus S_{Q-1}$. If (a), then $\Lambda_{R_{Q-1},R_Q} = 0$. If (b), then $\Lambda_{H_{Q-1}\setminus S_{Q-1},R_Q} = 0$. In either case, this leads to the block decomposition for the marginal precision matrix as

$$C^{-1}_{H_Q} = \begin{pmatrix} \Omega_{H_{Q-1}\setminus S_{Q-1}} & \Omega_{H_{Q-1}\setminus S_{Q-1},S_{Q-1}} & 0 \\ \Omega^T_{H_{Q-1}\setminus S_{Q-1},S_{Q-1}} & \Omega_{S_{Q-1}} & \Omega_{S_{Q-1},R_{Q-1}} \\ 0 & \Omega^T_{S_{Q-1},R_{Q-1}} & \Omega_{R_{Q-1}} \end{pmatrix}$$

that is the zero-structure of the overall graph is maintained for a subgraph where the nodes in $R_Q$ have been removed.

The same argument can be applied when removing successive residuals $R_{Q-1}$, $R_{Q-2}$ etc., since each time the removal results in another decomposable graph.

##### S.1.5 Derivation of posterior full conditionals for $\boldsymbol{\beta}$

###### Generic prior

Consider a general spike-and-slab prior for the regression coefficients, expressed in terms of its precision matrix $W$

$$\boldsymbol{\beta}|\boldsymbol{\gamma} \sim \boldsymbol{\gamma}\, \mathrm{N}\left(\mathbf{0}, W^{-1}\right) + \prod_{k,j} \left(1-\gamma_{kj}\right)\delta_0,$$

where $\boldsymbol{\beta} = \mathrm{vec}(\boldsymbol{\beta}_1, \ldots, \boldsymbol{\beta}_s)$ is the vector of the regression coefficients and $\boldsymbol{\gamma} = \mathrm{vec}(\boldsymbol{\gamma}_1, \ldots, \boldsymbol{\gamma}_s)$ is the vector of binary indicators. We notate the zero element $\delta_0$, the Dirach delta function in 0, and all non-zero elements (relative to non-zero $\boldsymbol{\gamma}$ coefficients) with $\boldsymbol{\beta}_{\boldsymbol{\gamma}}$ and, from here on, we will concern ourselves only with the slab part of the prior.

Its prior full conditional can straightforwardly be written as

$$\boldsymbol{\beta}_{\boldsymbol{\gamma}_k}|\boldsymbol{\beta}_{\backslash k} \sim \mathrm{N}\left((W_k)^{-1}W_{k,\backslash k}\boldsymbol{\beta}_{\backslash k}\ ,\ (W_k)^{-1}\right),$$

where the subscript "$\backslash k$" implies that the vector (or matrix) consists of all the elements except those that are related to the $k$-th response, $W_k$ is the $k$-th diagonal block with dimension $|\boldsymbol{\gamma}_k|\times|\boldsymbol{\gamma}_k|$ corresponding to regression coefficients for outcome $k$ and $W_{k,\backslash k}$ is its relative off-diagonal part.

Let's define

$$U^{(k-1)} = \Big((\boldsymbol{y}_1 - X_{\boldsymbol{\gamma}_1}\boldsymbol{\beta}_{\boldsymbol{\gamma}_1})\quad (\boldsymbol{y}_2 - X_{\boldsymbol{\gamma}_2}\boldsymbol{\beta}_{\boldsymbol{\gamma}_2})\quad \ldots\quad \big(\boldsymbol{y}_{k-1} - X_{\gamma_{k-1}}\boldsymbol{\beta}_{\boldsymbol{\gamma}_{k-1}}\big)\Big)\,.$$

For the calculations below, we need the indexing sets $\mathcal{L}(k), \mathcal{M}(k)$ and $\mathcal{H}(k,m)$, defined in Equation (7) of the main text. We obtain the posterior full conditional by isolating all the likelihood terms that depend on $\boldsymbol{\beta}_{\boldsymbol{\gamma}_k}$, which means not only the $k$-th term in the likelihood, but also all terms $m \in \mathcal{M}(k)$ where $\boldsymbol{\beta}_{\gamma_k}$ is found inside $U^{(m-1)}$, i.e.,

$$\begin{aligned}
\log p(\boldsymbol{\beta}_{\gamma_k}|Y, \boldsymbol{\beta}_{\setminus k}, \boldsymbol{\gamma}_k, \boldsymbol{\sigma}, \boldsymbol{\rho}, W) \propto& \\
&-\frac{1}{2}\left(\boldsymbol{\beta}_{\boldsymbol{\gamma}_k} + (W_k)^{-1} W_{k,-k} \boldsymbol{\beta}_{\setminus k}\right)^T W_k \left(\boldsymbol{\beta}_{\boldsymbol{\gamma}_k} + (W_k)^{-1} W_{k,-k} \boldsymbol{\beta}_{\setminus k}\right) \\
&-\frac{1}{2}\frac{1}{\sigma_k^2}\left(X_{\boldsymbol{\gamma}_k}\boldsymbol{\beta}_{\boldsymbol{\gamma}_k} - \left(\boldsymbol{y}_k - U^{(k-1)}\boldsymbol{\rho}_k\right)\right)^T \left(X_{\boldsymbol{\gamma}_k}\boldsymbol{\beta}_{\boldsymbol{\gamma}_k} - \left(\boldsymbol{y}_k - U^{(k-1)}\boldsymbol{\rho}_k\right)\right) \\
&\sum_{m\in\mathcal{M}(k)} -\frac{1}{2}\frac{1}{\sigma_m^2}\left(X_{\boldsymbol{\gamma}_m}\boldsymbol{\beta}_{\boldsymbol{\gamma}m} - \boldsymbol{y}_m + U^{(m-1)}\boldsymbol{\rho}_m\right)^T \left(X_{\boldsymbol{\gamma}_m}\boldsymbol{\beta}_{\boldsymbol{\gamma}m} - \boldsymbol{y}_m + U^{(m-1)}\boldsymbol{\rho}_m\right).
\end{aligned}$$

We can also write

$$U^{(m-1)}\boldsymbol{\rho}_m = \sum_{h\in\mathcal{H}(k,m)} \left(\boldsymbol{y}_h - X_{\boldsymbol{\gamma}_h}\boldsymbol{\beta}_{\boldsymbol{\gamma}_k}\right)\rho_{mh} + \boldsymbol{y}_k\rho_{mk} - X_{\boldsymbol{\gamma}_k}\boldsymbol{\beta}_{\boldsymbol{\gamma}_k}\rho_{mk},$$

thus

$$\begin{aligned}
&\propto \boldsymbol{\beta}_{\gamma_k}^T W_k \boldsymbol{\beta}_{\gamma_k} - 2\boldsymbol{\beta}_{\gamma_k}^T\left(-\left(W_k\right)^{-1} W_{k,\setminus k}\boldsymbol{\beta}_{\setminus k}\right) + \cdots + \\
&\frac{1}{\sigma_k^2}\left[\boldsymbol{\beta}_{\gamma_k}^T X_{\gamma_k}^t X_{\gamma_k}\boldsymbol{\beta}_{\gamma_k} - 2\boldsymbol{\beta}_{\gamma_k}^T X_{\gamma_k}^T\left(\boldsymbol{y}_k - \sum_{l\in\mathcal{L}(k)}\left(\boldsymbol{y}_j - X_{\gamma_l}\boldsymbol{\beta}_{\gamma_l}\right)\rho_{kl}\right) + \ldots\right] + \\
&\sum_{m\in\mathcal{M}(k)}\frac{1}{\sigma_m^2}\left[\rho_{mk}^2\boldsymbol{\beta}_{\gamma_k}^T X_{\gamma_k}^T X_{\gamma_k}\boldsymbol{\beta}_{\gamma_k} + \right. \\
&\quad\left. - 2\boldsymbol{\beta}_{\gamma_k}^T X_{\gamma_k}^T\rho_{mk}\left(\boldsymbol{y}_k\rho_{mk} - \left(\boldsymbol{y}_m - X_{\gamma_m}\boldsymbol{\beta}_{\gamma_m}\right) + \sum_{h\in\mathcal{H}(k,m)}\left(\boldsymbol{y}_h - X_{\gamma_h}\boldsymbol{\beta}_{\gamma_h}\right)\rho_{mh}\right) + \ldots\right]
\end{aligned}$$

$$\begin{aligned}
&\propto \boldsymbol{\beta}_{\gamma_k}^T\left(W_k + X_{\gamma_k}^t X_{\gamma_k}\left(\sigma_k^{-2} + \sum_{m\in\mathcal{M}(k)}\rho_{mk}^2\sigma_m^{-2}\right)\right)\boldsymbol{\beta}_{\gamma_k} \\
&- 2\boldsymbol{\beta}_{\gamma_k}^T\Bigg(\left(-\left(W_k\right)^{-1} W_{k,\setminus k}\boldsymbol{\beta}_{\setminus k}\right) + X_{\gamma_k}^T\Bigg[\sigma_k^{-2}\left(\boldsymbol{y}_k - \sum_{l\in\mathcal{L}(k)}\left(\boldsymbol{y}_j - X_{\gamma_l}\boldsymbol{\beta}_{\gamma_l}\right)\rho_{kl}\right) + \\
&\sum_{m\in\mathcal{M}(k)}\rho_{mk}\sigma_m^{-2}\left(\boldsymbol{y}_k\rho_{mk} - \left(\boldsymbol{y}_m - X_{\gamma_m}\boldsymbol{\beta}_{\gamma_m}\right) + \sum_{h\in\mathcal{H}(k,m)}\left(\boldsymbol{y}_h - X_{\gamma_h}\boldsymbol{\beta}_{\gamma_h}\right)\rho_{mh}\right)\Bigg]\Bigg),
\end{aligned}$$

where we omitted all the terms not dependent on $\boldsymbol{\beta}_{\gamma_k}$. It is also worth mentioning that $\lambda_k = \left(\sigma_k^{-2} + \sum_{m\in\mathcal{M}(k)} \rho_{mk}\sigma_m^{-2}\right)$, the $k$-th diagonal element of the precision matrix $\Lambda = C^{-1}$. From here, we find that

$$\boldsymbol{\beta}_{\gamma_k}|\boldsymbol{y}, \boldsymbol{\beta}_{\backslash k}, \boldsymbol{\gamma}_k, \boldsymbol{\sigma}^2, \boldsymbol{\rho}, W, \xi \sim \mathrm{N}\left((\widetilde{W}_k)^{-1}\left(X_{\gamma_k}^T\widetilde{\boldsymbol{y}}_k - (W_k)^{-1}W_{k,\backslash k}\boldsymbol{\beta}_{\backslash k}\right)\ , (\widetilde{W}_k)^{-1}\right)$$

with $\xi$ the perfect elimination ordering, $\widetilde{W}_k = X_{\gamma_k}^T X_{\gamma_k}\lambda_k + W_k$ and

$$\widetilde{\boldsymbol{y}}_k = \sigma_k^{-2}\left(\boldsymbol{y}_k - \sum_{l\in\mathcal{L}(k)}\left(\boldsymbol{y}_l - X_{\gamma_l}\beta_{\gamma_l}\right)\rho_{kl}\right) + \sum_{m\in\mathcal{M}(k)}\rho_{mk}\sigma_m^{-2}\left(\boldsymbol{y}_k\rho_{mk} - \left(\boldsymbol{y}_m - X_{\gamma_m}\boldsymbol{\beta}_{\gamma_m}\right) + \sum_{h\in\mathcal{H}(k,m)}\left(\boldsymbol{y}_h - X_{\gamma_h}\boldsymbol{\beta}_{\gamma_h}\right)\rho_{mh}\right).$$

These expressions are fairly general. By specifying the prior precision $W$ we will be able to obtain simpler and more useful expressions.

**Independent prior**

Consider for example the independent prior where

$$\boldsymbol{\beta}|\boldsymbol{\gamma} \sim \boldsymbol{\gamma}\,\mathrm{N}\left(\mathbf{0}, w^{-1}\mathbb{I}_{sp}\right) + \prod_{k,j}\left(1-\gamma_{kj}\right)\delta_0.$$

All the elements are independent and thus the prior full conditional for the $\boldsymbol{\beta}$ vector relative to one response will be

$$\boldsymbol{\beta}_{\gamma_k} \sim \mathrm{N}\left(\mathbf{0}, w^{-1}\mathbb{I}_{|\gamma_k|}\right).$$

Proceeding similarly as in the previous section, we obtain the following posterior full conditional

$$\boldsymbol{\beta}_{\gamma_k}|Y, \boldsymbol{\beta}_{\backslash k}, \boldsymbol{\gamma}_k, \boldsymbol{\sigma}^2, \boldsymbol{\rho}, w, \xi \sim \mathrm{N}\left((\widetilde{W}_k)^{-1}X_{\gamma_k}^T\widetilde{\boldsymbol{y}}_k\ , (\widetilde{W}_k)^{-1}\right)$$

with $\widetilde{W}_k = X_{\boldsymbol{\gamma}_k}^T X_{\boldsymbol{\gamma}_k} \lambda_k + \frac{1}{w}\mathbb{I}_{|\gamma_k|}$ and $\widetilde{\boldsymbol{y}}_k$ as defined above.

Here, the independence assumption implies that $W_{k,\setminus k}$ is $\mathbf{0}$ and thus the computations simplify considerably.

##### $g$-prior

Another case of interest is the $g$-prior (Zellner, 1986), for which we define

$$W^g = \frac{1}{w}\left(\mathcal{X}_{\boldsymbol{\gamma}}^T \left(C \otimes \mathbb{I}_n\right)^{-1} \mathcal{X}_{\boldsymbol{\gamma}}\right).$$

After some straightforward algebra, it is possible to see that in this case $W^g$ is a block-matrix with $\frac{\lambda_{kj}}{w} X_{\boldsymbol{\gamma}_k}^T X_{\boldsymbol{\gamma}_j}$ in its $k, j$ block, with $k, j \in 1, \ldots, s$. Thus, we have that $W_k^g$, the prior full conditional precision matrix, is $W_k^g = \frac{1}{w}\left(X_{\boldsymbol{\gamma}_k}^T X_{\boldsymbol{\gamma}_k} \lambda_k\right)$ and $W_{k,\setminus k}^g = \frac{1}{w} X_{\boldsymbol{\gamma}_k}^T A$ with

$$A = \begin{pmatrix} & \cdots & \\ \cdots & X_{\gamma_j}\lambda_{k,j} & \cdots \\ & \cdots & \end{pmatrix}_{j \in \{1,\ldots,s\}\setminus k}.$$

Hence, the posterior full conditional simplifies to

$$\boldsymbol{\beta}_{\gamma_k} | Y, \boldsymbol{\beta}_{\setminus k}, \boldsymbol{\gamma}_k, \boldsymbol{\sigma}^2, \boldsymbol{\rho}, w, \xi \sim \mathrm{N}\left( (\widetilde{W}_k^g)^{-1} X_{\boldsymbol{\gamma}_k}^T \left( \widetilde{\boldsymbol{y}}_k - \frac{1}{w} A \boldsymbol{\beta}_{\setminus k} \right) \; , \; (\widetilde{W}_k^g)^{-1} \right)$$

with $\widetilde{W}_k^g = \frac{w+1}{w}\left(X_{\boldsymbol{\gamma}_k}^T X_{\boldsymbol{\gamma}_k} \lambda_k\right)$ and $\widetilde{\boldsymbol{y}}_k$ as defined above.

#### S.2 Supplementary Figures

Here, we present some additional graphs and figures of the simulation study and the analysis of the NFBC66 data set presented in Section 4 and Section 5 of the main text, respectively. See figures captions and main text for a more in-depth description.

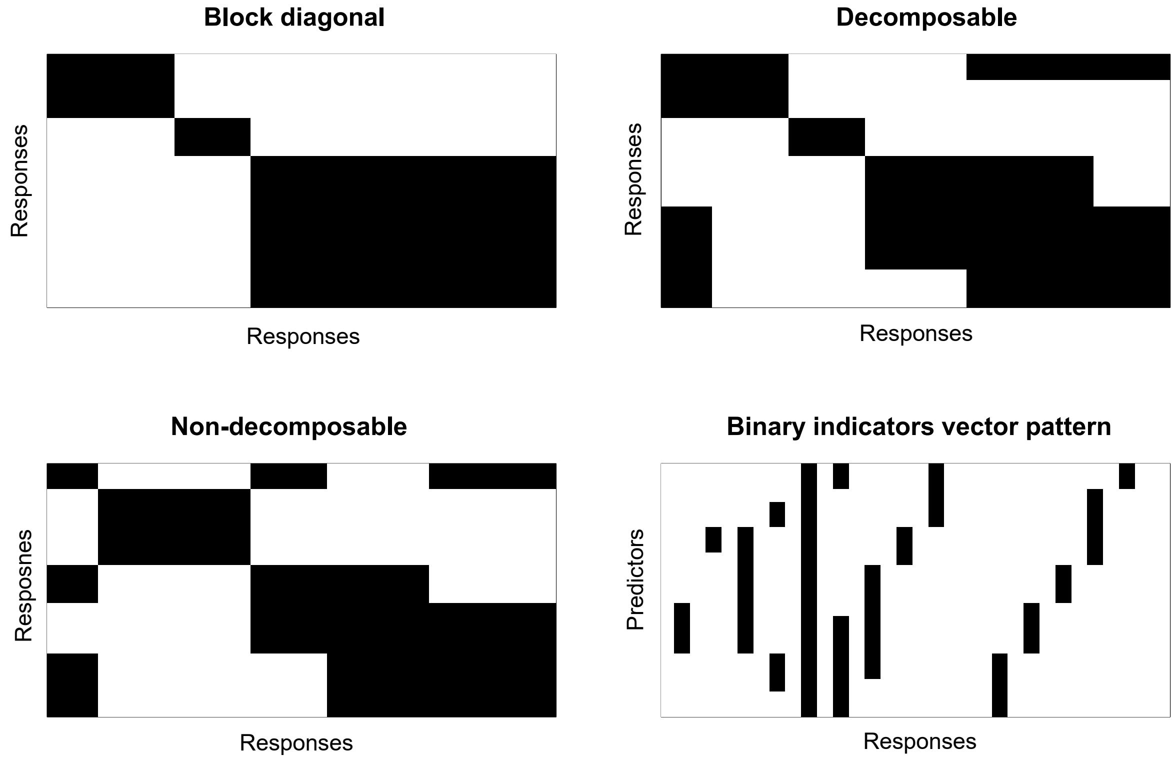


Figure S.2: Top-left, top-right and bottom-left panels: Graph structures, represented as adjacency matrices, used in the simulation study presented in Section 4.1 of the main text (black tassels represent edges). Bottom-right panel: Example of the simulated pattern of the binary indicators vector $\boldsymbol{\gamma}$ for $G =$ non-decomposable graphical model with responses on the rows and predictors on the columns (black tassels represent associations).

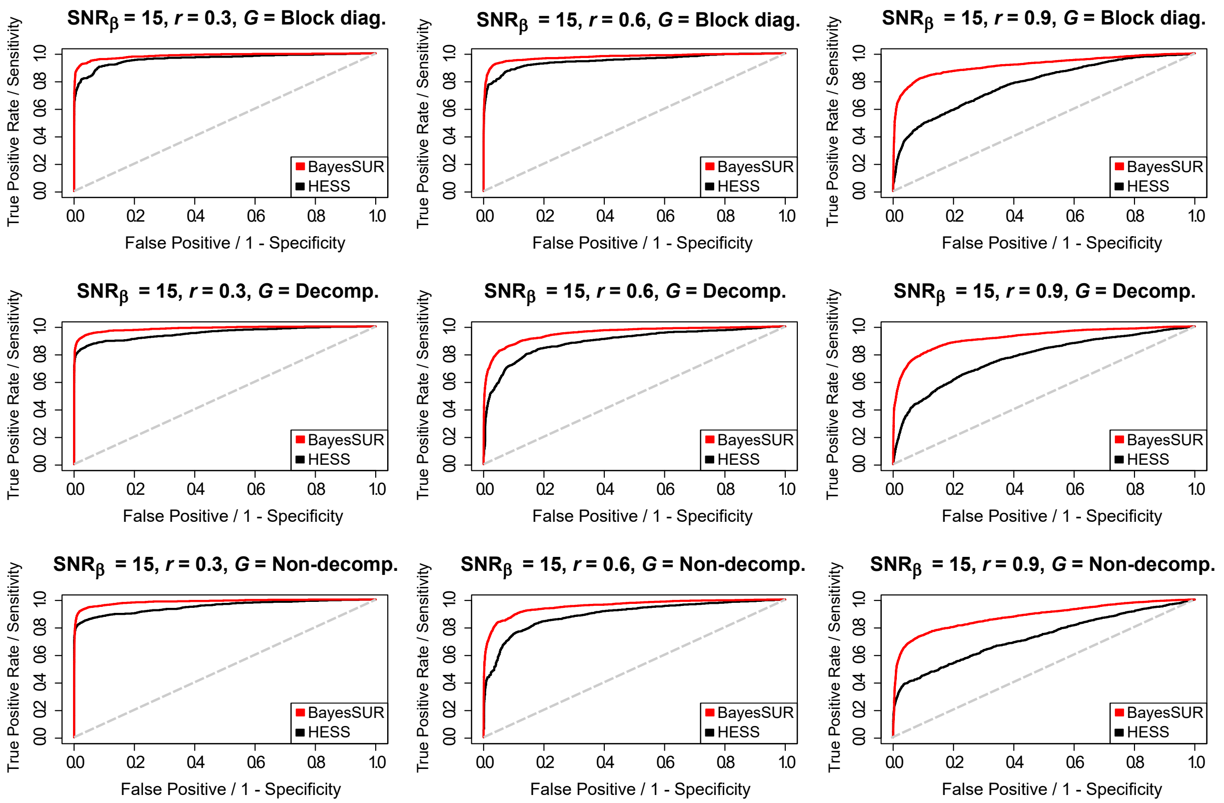


Figure S.3: Averaged (over 20 simulated replicates) ROC curves for BayesSUR with covariance selection (red line) and HESS (black line) to compare the variable selection performance of the two methods for different combinations of the simulated graphical model $G$ and the (residual) correlation between responses $r$ and with signal-to-noise ratio for the predictors $\text{SNR}_\beta = 15$ as described in Section 4.1 of the main text.

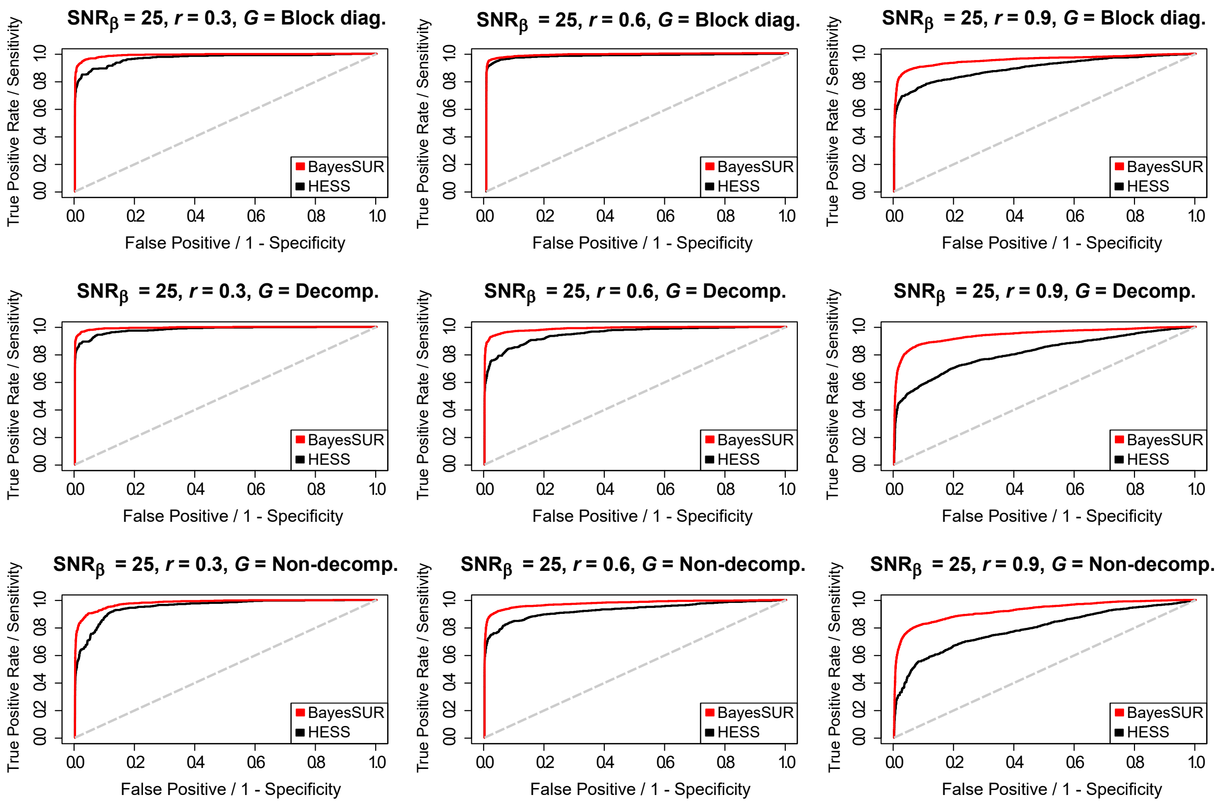


Figure S.4: Averaged (over 20 simulated replicates) ROC curves for BayesSUR with covariance selection (red line) and HESS (black line) to compare the variable selection performance of the two methods for different combinations of the simulated graphical model $G$ and the (residual) correlation between responses $r$ and with signal-to-noise ratio for the predictors $\text{SNR}_\beta = 25$ as described in Section 4.1 of the main text.

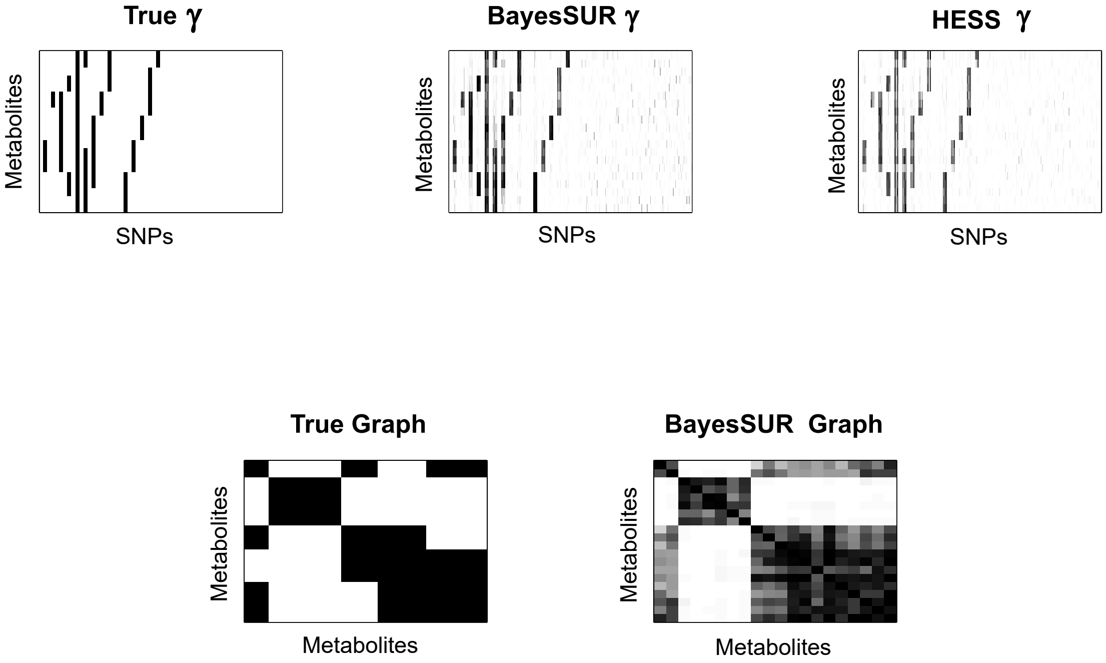


Figure S.5: Results based on the analysis of one simulated data set generated with $\mathrm{SNR}_\beta = 15$, $G$ = non-decomposable graphical model and (residual) correlation between responses $r = 0.6$ and presented in Section 4.1 of the main text. Top panels: True and estimated marginal posterior inclusion probabilities for both BayesSUR with covariance selection and HESS. HESS estimation appears quite good even for this intermediate level of $\mathrm{SNR}_\beta$, but the recovered signal for the simulated associations is noticeable less strong than BayesSUR. Bottom panels: True and estimate posterior marginal edge inclusion probabilities used to reconstruct the graph, where the BayesSUR algorithm seems to converge to a denser triangulations of the true graph.

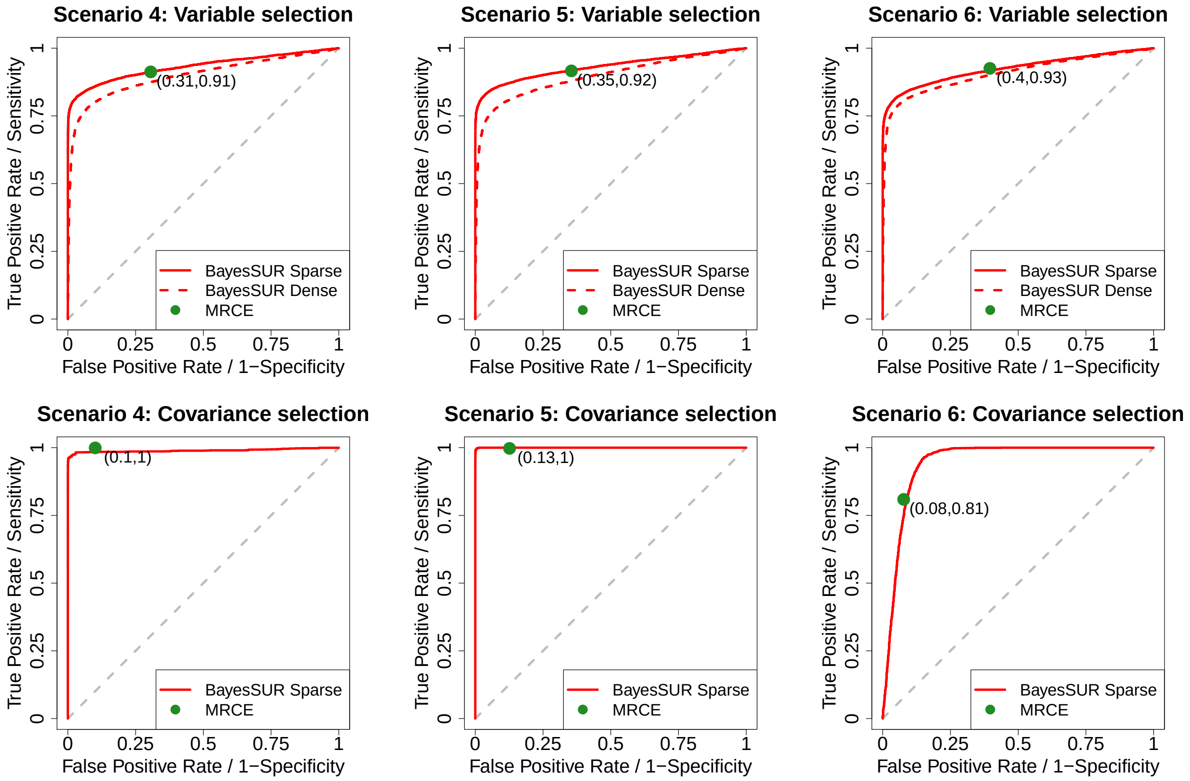


Figure S.6: Averaged (over 20 simulated replicates) ROC curves to compare the selection performance of the non-zero regression coefficients (top panels) and non-zero elements of the precision matrix (bottom panels) for the methods considered: BayesSUR with covariance selection (solid red line), BayesSUR with dense covariance estimation (dashed red line) and MRCE (green dot). For MRCE, each dot represents the averaged specificity and sensitivity of the corresponding penalised likelihood solution. As described in Section 4.2 of the main text, different scenarios are obtained by specifying distinct Toeplitz matrices for the inverse error covariance. Given that the computational time of SSUR Direct and SSUR Indirect for each simulated replicate is greater than the High Performance Computer cluster time wall (36h), results are not reported. BayesSUR with covariance selection and MRCE perform similarly in terms of sensitivity, but MRCE seems to include more false positives. When a large number of responses is considered ($s = 150$), the estimation of the residual covariance structure seems paramount for the effective detection of important predictors, as shown in the comparison between the two instances of the BayesSUR algorithm.

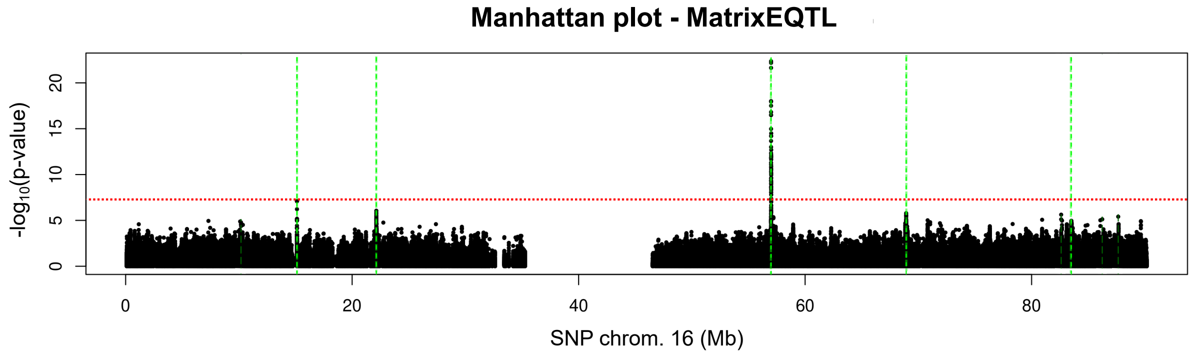


Figure S.7: Manhattan plot of -log transformed p-values in chromosome 16 obtained from the `MatrixEQTL` package (Shabalin, 2012) for the NFBC66 data set. The red dotted horizontal line represents the standard $5 \times 10^{-8}$ genome-wide selection cut-off and green dashed vertical dashed lines highlight the association found by BayesSUR with covariance selection, as reported in Section 5 of the main text.

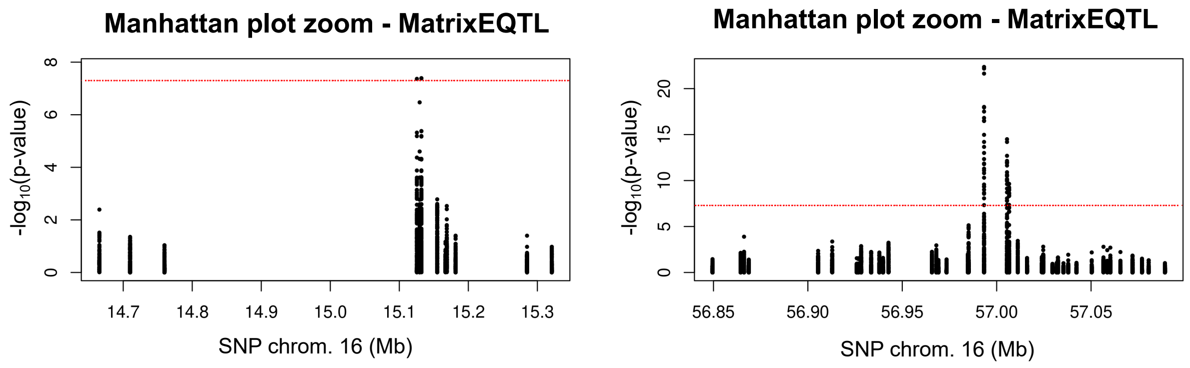


Figure S.8: Manhattan plots of -log transformed p-values in two smaller regions of chromosome 16 obtained from the `MatrixEQTL` package (Shabalin, 2012) for the NFBC66 data set. Red dotted horizontal lines represent the standard $5 \times 10^{-8}$ genome-wide selection cut-off. Multiple close SNPs are deemed predictive, reaffirming the relevance of the locus but making difficult to specifically point to important variants.

#### S.3 Supplementary Tables

Here, we present some additional tables of the simulation study and the analysis of the NFBC66 data set presented in Section 4 and Section 5 of the main text, respectively. See tables captions and main text for a more in-depth description.

Table S.1: Averaged (over 20 simulated replicates) true positive and false positive rates for the graph estimation after thresholding at 0.5 the posterior marginal edge inclusion probabilities. Results are reported for $\text{SNR}_\beta = 15$ and $\text{SNR}_\beta = 25$ and for different combinations of the graphical model $G$ and the (residual) correlation between responses $r$.

| $\text{SNR}_\beta$ | $G$ | $r$ | True positive rate | False positive rate |
|---|---|---|---|---|
| 15 | Block diagonal | 0.3 | 0.954 | 0.005 |
| | Block diagonal | 0.6 | 0.953 | 0 |
| | Block diagonal | 0.9 | 0.987 | 0.021 |
| | Decomposable | 0.3 | 0.931 | 0.071 |
| | Decomposable | 0.6 | 0.979 | 0.080 |
| | Decomposable | 0.9 | 0.981 | 0.122 |
| | Non-decomposable | 0.3 | 0.968 | 0.070 |
| | Non-decomposable | 0.6 | 0.990 | 0.170 |
| | Non-decomposable | 0.9 | 0.993 | 0.110 |
| 25 | Block diagonal | 0.3 | 0.953 | 0.003 |
| | Block diagonal | 0.6 | 0.975 | 0 |
| | Block diagonal | 0.9 | 0.993 | 0 |
| | Decomposable | 0.3 | 0.960 | 0.084 |
| | Decomposable | 0.6 | 0.979 | 0.092 |
| | Decomposable | 0.9 | 0.978 | 0.150 |
| | Non-decomposable | 0.3 | 0.935 | 0.110 |
| | Non-decomposable | 0.6 | 0.976 | 0.141 |
| | Non-decomposable | 0.9 | 1 | 0.188 |

Table S.2: Averaged (over 20 simulated replicates) computational time in minutes for the algorithms considered: BayesSUR with covariance selection (Sparse), BayesSUR with dense covariance estimation (Dense) and MRCE with different numbers of candidate values for the penalty parameters where the cross-validation procedure is performed. Given that the computational time of SSUR Direct and SSUR Indirect for each simulated replicate is greater than the High Performance Computer cluster time wall (36h), results are not reported. For the same reason, we do not report the computational time for the MRCE algorithm when 40 and 200 candidate values for the penalty parameters $\lambda_1$ and $\lambda_2$ are considered.

| Algorithm | | Scenario 4 | Scenario 5 | Scenario 6 |
|---|---|---|---|---|
| BayesSUR | Sparse | $1,338$ | $1,404$ | $1,412$ |
| | Dense | $1,761$ | $1,821$ | $1,841$ |
| MRCE | 200 candidate values for $\lambda_1$ and $\lambda_2$ | - | - | - |
| | 40 candidate values for $\lambda_1$ and $\lambda_2$ | - | - | - |
| | 4 candidate values for $\lambda_1$ and $\lambda_2$ | 997 | 229 | 229 |

Table S.3: Uncovered hotspots genetic variants and associated metabolites after thresholding the marginal posterior inclusion probabilities at 0.5.

| SNP | Associated metabolites | | | | |
|---|---|---|---|---|---|
| rs4985124 | FAw3.FA | FAw6.FA | | | |
| rs7191766 | L.LDL.P | L.LDL.L | L.LDL.PL | L.LDL.C | L.LDL.CE |
| | L.LDL.FC | M.LDL.P | M.LDL.L | M.LDL.PL | M.LDL.C |
| | M.LDL.CE | M.LDL.FC | S.LDL.P | S.LDL.L | S.LDL.PL |
| | S.LDL.C | S.LDL.CE | S.LDL.FC | S.HDL.C | S.HDL.CE |
| | Serum.C | LDL.C | EstC | FreeC | LA |
| | FAw3 | FAw6 | PUFA | | |
| rs3764261 | XXL.VLDL.C | XXL.VLDL.CE | XL.VLDL.CE | L.VLDL.CE | M.VLDL.CE |
| | S.VLDL.PL | S.VLDL.C | S.VLDL.CE | XS.VLDL.C | XS.VLDL.CE |
| | XL.HDL.P | XL.HDL.L | XL.HDL.PL | XL.HDL.C | XL.HDL.CE |
| | XL.HDL.FC | L.HDL.P | L.HDL.L | L.HDL.PL | L.HDL.C |
| | L.HDL.CE | L.HDL.FC | M.HDL.P | M.HDL.L | M.HDL.PL |
| | M.HDL.C | M.HDL.CE | M.HDL.FC | M.HDL.TG | S.HDL.TG |
| | LDL.D | HDL.D | VLDL.C | HDL.C | HDL2.C |
| | HDL3.C | TotPG | TG.PG | PC | SM |
| | TotChol | ApoA1 | ApoB | LA | LA.FA |
| rs12102766 | XXL.VLDL.PL | XXL.VLDL.C | XXL.VLDL.CE | XXL.VLDL.FC | XL.VLDL.P |
| | XL.VLDL.L | XL.VLDL.PL | XL.VLDL.C | XL.VLDL.CE | XL.VLDL.FC |
| | L.VLDL.P | L.VLDL.L | L.VLDL.PL | L.VLDL.C | L.VLDL.CE |
| | L.VLDL.FC | M.VLDL.L | M.VLDL.PL | M.VLDL.C | M.VLDL.CE |
| | M.VLDL.FC | VLDL.C | | | |
| rs931406 | L.LDL.PL | M.LDL.PL | S.LDL.P | S.LDL.L | S.LDL.PL |
| | Serum.C | EstC | FreeC | TotPG | SFA |

Table S.4: Relevant associations for non-hotspot SNPs after thresholding the marginal posterior inclusion probabilities at 0.5.

| SNP | Associated Metabolite |
|---|---|
| rs17683096 | LDL.D |
| rs4782717 | XL.HDL.CE |
| rs4843914 | Cit |
| rs2544630 | Gln |

### References


Carvalho, C. M., H. Massam, and M. West (2007). Simulation of hyper-inverse Wishart distributions in graphical models. *Biometrika 94*, 647–659.

Lauritzen, S. L. (1996). *Graphical Models*. Oxford University Press, New York.

Paulsen, V. I., S. C. Power, and R. R. Smith (1989). Schur products and matrix completions. *J. Funct. Anal. 85*(1), 151–178.

Shabalin, A. (2012). Matrix eQTL: Ultra fast eQTL analysis via large matrix operations. *Bioinformatics 28*(10), 1353–1358.

Zellner, A. (1986). On assessing prior distributions and Bayesian regression analysis with $g$-prior distributions. In P. Goel and A. Zellner (Eds.), *Bayesian Inference and Decision Techniques: Essays in Honor of Bruno de Finetti*, Volume 6, pp. 233–243. Elsevier Science Publishers, Inc., New York.
